# On the Edge of Empire: Paleogenomic Insights into Roman Dacia

**DOI:** 10.64898/2026.04.18.719386

**Authors:** Flavio De Angelis, Ileana Buzic, Kalina Kassadjikova, Adrian Cosmin Bolog, Anca Timofan, John Pearce, Mihai Gligor, Lars Fehren-Schmitz, Carlos Eduardo G. Amorim

**Author notes:** **Correspondence:** (F.D.A.), (C.E.G.A.).

## Abstract

The Roman province of Dacia, located north of the Danube frontier, represented a key zone of cultural and demographic interaction during the Imperial period. However, the biological impact of Roman colonization in this region has not been characterized using genomic data. Here, we analyze genome-wide data from 34 individuals recovered from the Apulum-*Dealul Furcilor* necropolis, one of the largest funerary complexes in Roman Dacia. The genome-wide data reveal pronounced genetic heterogeneity within this population, reflecting its position at the intersection of Eastern Europe, the Mediterranean, and West Asia. Notably, we observe a sex-biased pattern of ancestry. Female individuals show stronger affinities to Eastern European, Steppe, and Caucasus-associated populations, suggesting the persistence of local or regionally connected genetic lineages. In contrast, male individuals display closer genetic relationships with Mediterranean and North African groups, including populations associated with Roman and Punic contexts, indicating male-mediated gene flow linked to long-distance mobility. These findings highlight the complex demographic processes shaping Roman frontier communities, where local and incoming populations were integrated through asymmetric social dynamics. Our results provide genomic evidence consistent with sex-biased admixture in Roman Dacia and underscore the role of frontier regions as hubs of genetic and cultural interaction within the Roman Empire.

## Introduction

Starting around 500 BCE, the Roman Empire expanded gradually, eventually establishing a frontier stretching approximately 7,500 kilometers across Europe and North Africa by the 2^nd^ century CE, forming one of the most extensive territorial systems in ancient history^1^. Among its frontiers, the province of Dacia (Figure 1), located entirely north of the Danube River, emerged as a strategically vital region, serving as a critical defensive and administrative outpost for nearly 170 years^2^. The territory of Dacia, situated at a zone of transcontinental intersection, represents a unique melting pot where diverse cultural influences converged^3^. To the east, the Scythians contributed martial traditions, metalworking expertise, and trade networks spanning the Eurasian steppe^4^, while Greek colonies along the Black Sea coast introduced artistic, architectural, and political traditions^5^. These long-standing interactions shaped Dacia’s complex historical trajectory and formed the backdrop to its eventual incorporation into the Roman Empire in 106 CE following violent conquest led by the emperor Trajan (see Supplementary Note S1 for historical background).

**Figure 1:**
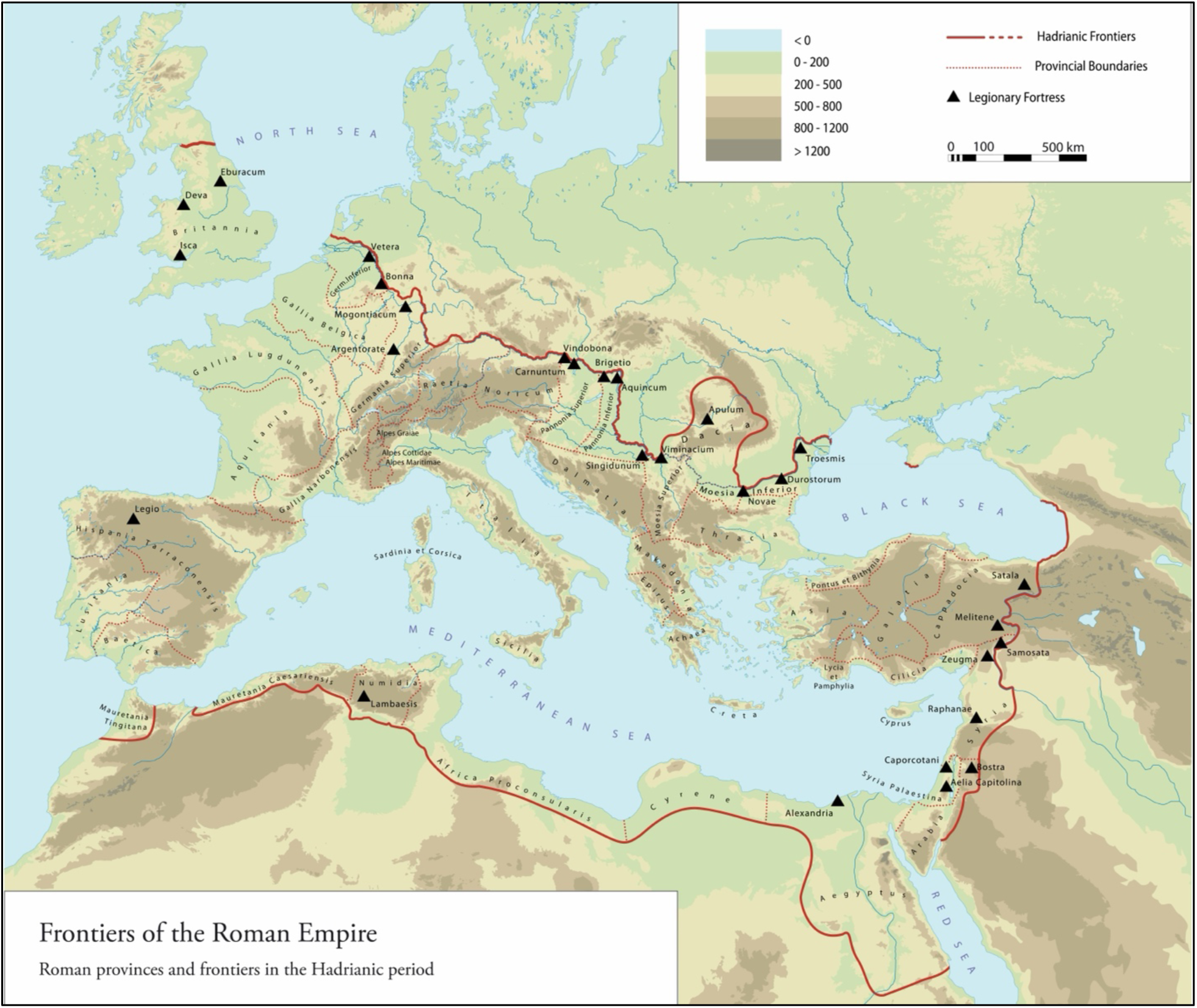
Location of Dacia in the context of Roman Empire. From *Danube Limes – UNESCO World Heritage* (Pen & Sword; CHC, University of Salzburg), by David Breeze and Kurt Schaller. Reproduced with permission of ao. Univ.-Prof. i.R. Dr. Andreas Schwarcz.

Through the analysis of ancient DNA (aDNA), this study aims to characterize the genetic makeup of the populations that inhabited Roman Dacia, shedding light on the extent of Roman colonization of local communities, and the influence of neighboring nomadic groups such as the Sarmatians. In doing so, we address key questions about how the Roman conquest and subsequent colonization shaped the genetic structure of the region, and to what extent the local population admixed with Roman settlers and other groups, in particular the Sarmatians^6^. More broadly, our work examines the impact of Roman rule on the genetic diversity of the province, exploring how the influx of Roman settlers, veterans, and civilians contributed to the patterns observed in this community, while also considering the long-term genetic legacy of Roman Dacia and investigating whether the genomic signatures of this period persisted in the region after the Roman withdrawal in 271 CE. By combining genomic data with historical and archaeological insights, this study aims to contribute to a comprehensive understanding of the demographic and genetic processes that shaped Roman Dacia. It seeks to bridge the gap between the biological and cultural dimensions of the province’s history, offering new perspectives on the interplay between Roman imperial policies, local populations, and neighboring groups in shaping the genomic landscape of this frontier region.

## Results

### Genomic data

Shotgun sequencing of 34 human DNA samples from the Alba Iulia-*Dealul Furcilor* (ADF) necropolis yielded ancient genomes with an average coverage ranging from ∼0.0001 to 0.604x (mean ∼0.1x). Contamination was estimated using both X-chromosome data (for male individuals) and mitochondrial DNA (for all individuals), confirming that all genomes included in downstream analyses had contamination rates <5%. Post-mortem deamination patterns confirmed the authenticity of the ancient DNA material (Supplementary Table S1). The number of SNPs detected from the 1240k panel (total of 1,233,013 biallelic variants) ranged from 86 to 515,068 across individuals; while the SNP sharing with the combined 1240k_HO dataset (total of 597,573 biallelic variants) ranged from 43 to 280,421 across individuals. A threshold of a minimum of 10,000 SNPs shared with the 1240k_HO set was applied to select genomes for downstream population genetic analysis, resulting in a final set of 14 individuals. Relatedness analysis performed on this subset revealed no close genetic relationships among individuals with sufficient SNP overlap (Supplementary Table S2).

### A population at the crossroads of Eurasia

Principal Component Analysis (PCA) placed the ADF individuals within a broad genetic space that bridges Eastern Europe, the Mediterranean, and West Asia (Figure 2). Rather than forming a cohesive cluster, ADF individuals are dispersed in the PCA space, suggesting a genetically diverse community. One individual (ADF_27) plotted closely with Western Asian populations, while the others were scattered among clusters representing Europeans, Mediterraneans, North Africans, and populations at the interface of Eastern Europe and West Asia. This heterogeneity suggests that the ADF population was not an isolated group, but rather a genetically diverse community situated within a contact zone. The observed pattern may reflect complex interactions, likely shaped by multiple waves of migration and admixture over time.

**Figure 2:**
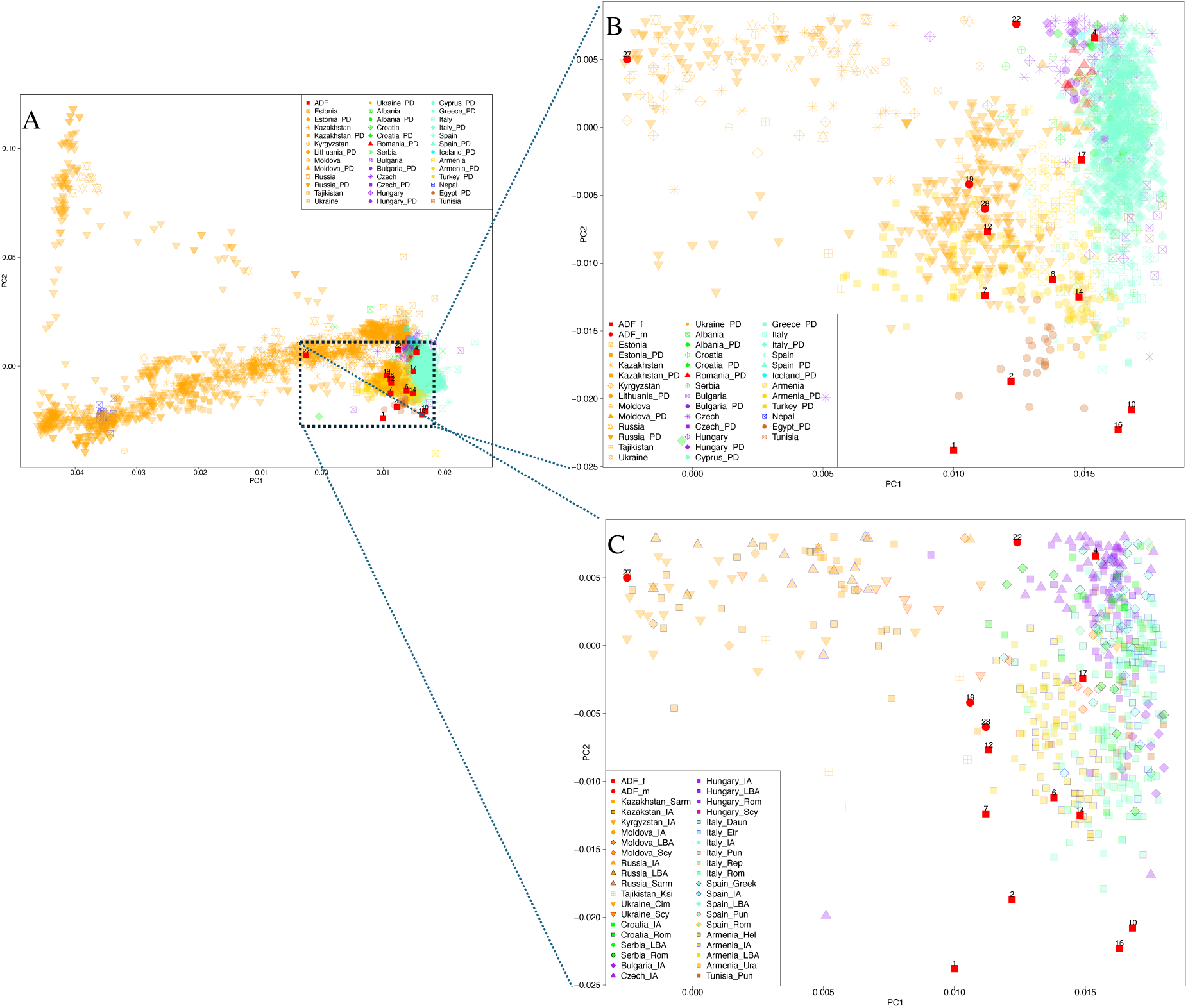
Principal component analysis of ancient and present-day populations highlighting the genetic structure of the ADF individuals. (**A**) Principal component analysis (PCA) based on genome-wide variation in present-day Eurasian and North African populations, with ancient individuals projected onto this space. (**B**) Zoomed-in view of the PCA space showing the distribution of ADF individuals relative to key reference populations, labeled by geographic origin and chronological attribution as Present Day (PD) or ancient individuals. (**C**) Zoomed-in view of the PCA space showing the distribution of ADF and other ancient individuals labeled by cultural attribution. Cim: Cimmerian; Daun: Daunian; Etr: Etruscan; Hel: Hellenistic; IA: Iron Age; Ksi: Ksiron; LBA: Late Bronze Age; Pun: Punic; Rep: Republican; Rom: Roman; Sarm: Sarmatian; Scy: Scythian; Ura: Urartian.

### Female ancestry: Steppe and Caucasus affinities

The ancestry profiles of ADF individuals, derived from an unsupervised clustering analysis, reveal pronounced sex-specific differences in ancestry composition (Figure 3, Supplementary Figure 1). In ADF_f (female) individuals, the predominant genetic contribution comes from Component 2 (green), accounting for 49.2% of their ancestry. This is followed by Component 1 (red) at 35.8% and a smaller proportion of Component 3 (blue) at 15.0%. In contrast, ADF_m (male) individuals exhibit a markedly different pattern: their ancestry is dominated by Component 1 (49.0%), followed by a substantial presence of Component 3 (30.4%), and a weaker representation of Component 2 (20.5%). These divergent profiles suggest that the male and female individuals in the ADF population may have drawn ancestry from at least partially distinct gene pools, likely shaped by sex-biased migration patterns, sex-biased population replacements, patrilocality, or other sociocultural and demographic processes reflecting the complex dynamics of human mobility and integration in the region.

**Figure 3:**
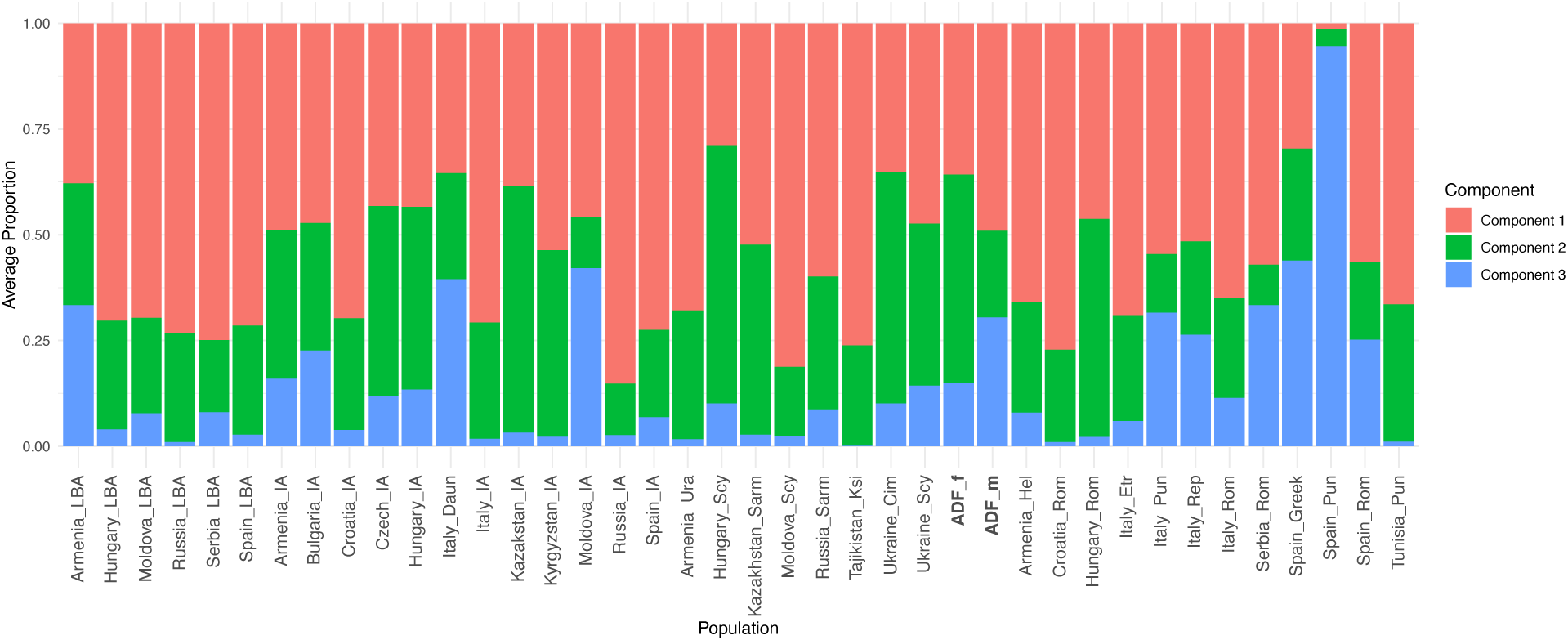
Ancestry profiles of ADF individuals in the context of ancient Eurasian populations. Unsupervised admixture analysis showing ancestry components inferred at *K* = 3 ancestral components. Each vertical bar represents an individual, and colors correspond to distinct ancestral components shared across ancient populations. Cim: Cimmerian; Daun: Daunian; Etr: Etruscan; Hel: Hellenistic; IA: Iron Age; Ksi: Ksiron; LBA: Late Bronze Age; Pun: Punic; Rep: Republican; Rom: Roman; Sarm: Sarmatian; Scy: Scythian; Ura: Urartian.

To contextualize the ancestry components observed in ADF_f individuals, we consider their genetic profiles in relation to a broad range of Iron Age and Late Bronze Age (LBA) populations. The dominant Component 2 in females shows strong parallels with ancient populations from the Eurasian Steppe and its peripheries, particularly during the early 1^st^ millennium BCE. For instance, this component constituted 60.8% of the genetic makeup in *Hungary_Scy*, 58.2% in *Kazakhstan_IA*, and 54.6% in *Ukraine_Cim*, reflecting the legacy of mobile pastoralist groups that expanded across the Steppe during the LBA-Iron Age transition.

These affinities were corroborated by outgroup f₃-statistics (Figure 4), in the form of f₃(ADF_f, X; YRI), revealing the highest levels of shared genetic drift with *Armenia_Ura* (0.111) and *Moldova_LBA* (0.111), populations dating to the 13^th^-9^th^ centuries BCE, as well as *Russia_LBA* (0.110) and *Tajikistan_Ksi* (0.111). Additional affinities with *Italy_IA* (0.109) and groups from the Carpathian Basin, including *Hungary_IA*, *Hungary_Scy*, and *Hungary_LBA* (0.105-0.108), suggest continued genetic interactions between the eastern and central parts of Europe throughout the Iron Age, likely facilitated by long-distance mobility and shifting social networks. Central European groups such as *Czech_IA* and *Croatia_IA* (∼0.106-0.108) further illustrate the geographic breadth of this shared ancestry.

**Figure 4:**
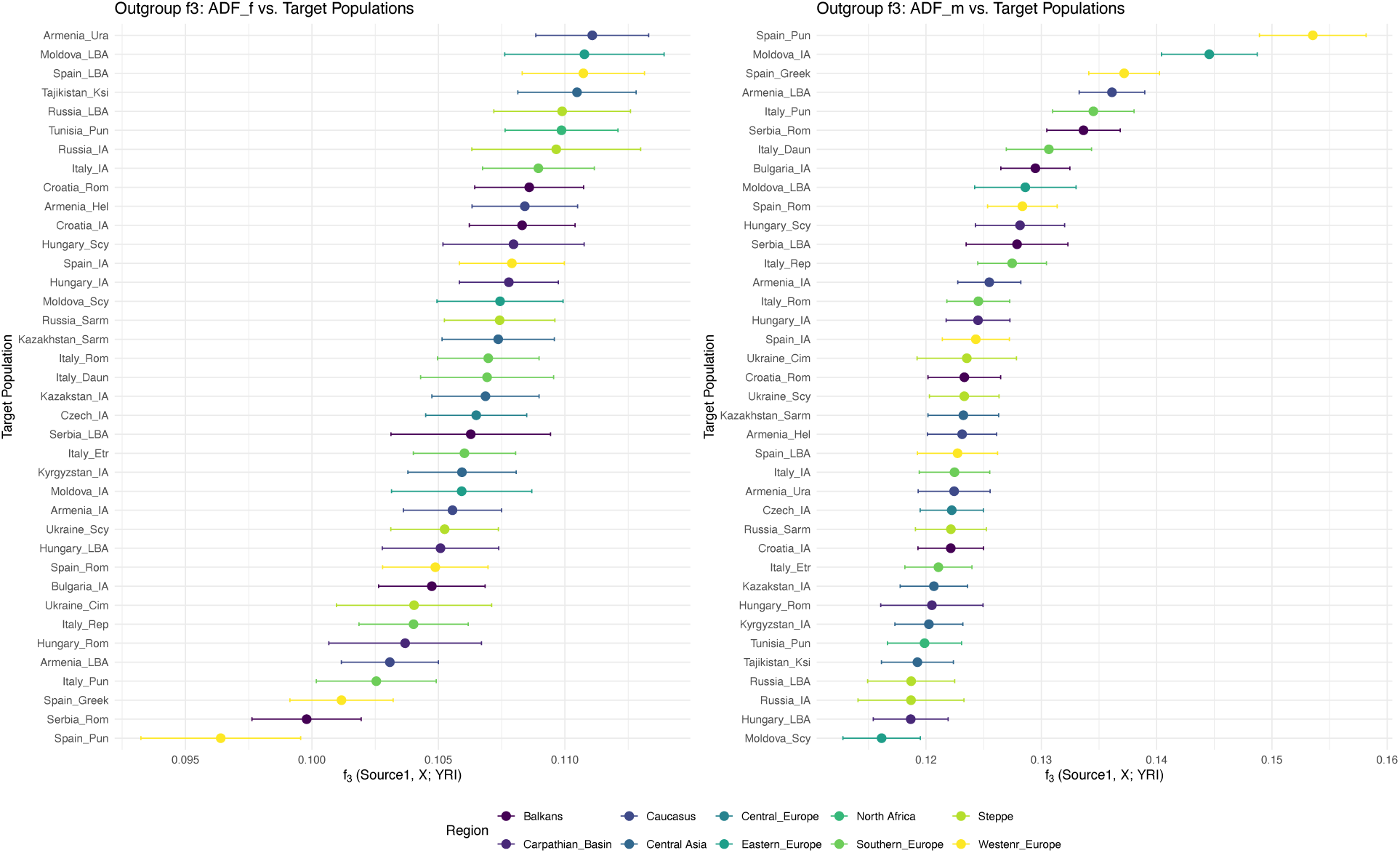
Outgroup f₃-statistics showing shared genetic drift between ADF individuals and ancient populations. Outgroup f₃-statistics of the form f₃(ADF_f, X; YRI) (left) and f₃(ADF_m, X; YRI) (right), measuring shared genetic drift between ADF individuals and a range of ancient Eurasian populations, using Yoruba (YRI) as an outgroup. Colors refer to different geographical regions. Cim: Cimmerian; Daun: Daunian; Etr: Etruscan; Hel: Hellenistic; IA: Iron Age; Ksi: Ksiron; LBA: Late Bronze Age; Pun: Punic; Rep: Republican; Rom: Roman; Sarm: Sarmatian; Scy: Scythian; Ura: Urartian.

These values indicate substantial genetic continuity between ADF_f individuals and Steppe-Caucasus-Danubian populations active between the 14^th^ and 4^th^ centuries BCE, suggesting either recent female-mediated gene flow from the East into the ADF community or alternatively a scenario in which female individuals were the sole surviving descendants of a local population with long-standing genetic ties to Steppe-Caucasus-Danubian groups, predating the Roman conquest of Dacia in the 2^nd^ century CE.

To further explore the temporal dimension of these affinities, we used D-statistics to assess shared ancestry between ADF_f and Iron Age populations. We observe some of the most significant signals in comparisons such as D(YRI, *Croatia_IA*; ADF_f, *Italy_IA*) with a Z-score of −20.122; D(YRI, *Armenia_Hel*; ADF_f, *Croatia_IA*) at −19.830; and D(YRI, *Tunisia_Pun*; ADF_f, *Croatia_Rom*) at −19.484. These results underscore deep genetic connections established during the LBA, sustained through the Iron Age, and preserved in ADF females, likely shaped by centuries of trans-regional movement and integration involving Steppe and Caucasus populations (Figure 5).

**Figure 5:**
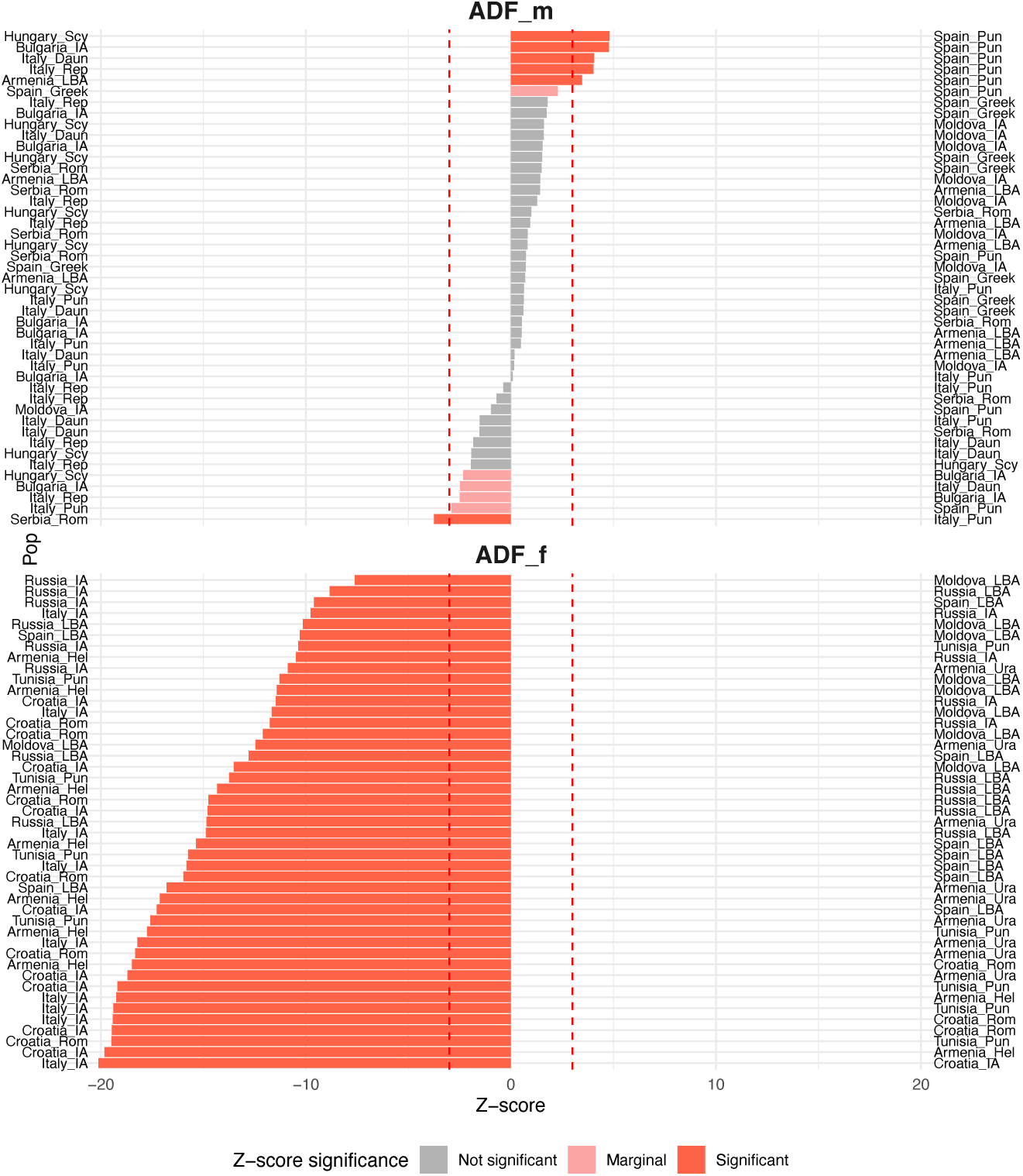
D-statistics highlighting differential genetic affinities of ADF females and males. D-statistics of the form D(ADF_f, X ; Y, YRI) showing allele sharing differences between ADF males (ADF_m, top panel) and ADF females (ADF_f, bottom panel), relative to a range of ancient populations (X, Y), using Yoruba (YRI) as an outgroup. Color bars mark significance thresholds (grey: not significant, with |Z| :< 2.5; pink: marginally significant, with 2.5 ≥ |Z| ≥ 3; red: significant, with |Z| ≥ 3). Cim: Cimmerian; Daun: Daunian; Etr: Etruscan; Hel: Hellenistic; IA: Iron Age; Ksi: Ksiron; LBA: Late Bronze Age; Pun: Punic; Rep: Republican; Rom: Roman; Sarm: Sarmatian; Scy: Scythian; Ura: Urartian.

### Male ancestry: A Mediterranean mosaic

In contrast to the female fraction, the ancestry of ADF_m (male) individuals reveals a markedly different profile (Figure 3), pointing to genetic contributions rooted in Mediterranean and North African maritime networks that were particularly active from the Late Bronze Age through the Classical and Roman periods, i.e. from the late 2^nd^ millennium BCE through the first centuries CE. The dominant admixture components in ADF_m – Component 1 and Component 3 – are strongly represented in Iron Age populations from the central and western Mediterranean. Specifically, *Italy_IA* and *Italy_Etr* exhibited Component 1 at 70.8% and 69.0%, respectively, while *Croatia_Rom* showed an even higher value (77.2%), highlighting shared ancestry with Romanized Balkan populations. In parallel, Component 3 was especially dominant in *Spain_Pun* (94.6%), and was also a major contributor in *Tunisia_Pun* and *Italy_Pun*, suggesting a strong genetic link between ADF_m individuals and Punic or North African populations engaged in Mediterranean trade and colonization.

These affinities were quantitatively confirmed through outgroup f₃-statistics, where ADF_m showed even higher values than ADF_f (Figure 4). The strongest signal in the dataset was observed with *Spain_Pun* (0.154), followed by *Italy_Pun* (0.135), *Serbia_Roman* (0.134), and *Bulgaria_IA* (0.129). Importantly, high f₃ values were also detected with *Armenia_LBA* (0.136) and *Moldova_IA* (0.145), reflecting deep-rooted Caucasus-Balkan connections for males, albeit embedded within broader Mediterranean patterns. These results are consistent with a scenario where ADF_m individuals were the product of male-mediated gene flow from western and southern populations, potentially linked to the movement of soldiers, sailors, merchants, or administrators during the Roman imperial expansion and, possibly, with earlier Phoenician mobility across the Mediterranean in the 1^st^ millennium BCE.

D-statistics supported these conclusions with strong and statistically significant patterns of allele sharing. For instance, D(YRI, *Spain_Pun*; ADF_m, *Hungary_Scy*) showed a Z-score of 4.798, and D(YRI, *Spain_Pun*; ADF_m, *Bulgaria_IA*) of 4.781, highlighting ADF_m’s close affinity to Punic-influenced western Mediterranean populations. Intriguingly, subtle genetic structure within Punic and Roman populations was also detected. For example, D(YRI, *Italy_Pun*; ADF_m, *Serbia_Rom*) yielded a significant negative value (Z = −3.754), while D(YRI, *Spain_Pun*; ADF_m, *Italy_Pun*) yielded only a marginally significant Z-score of 2.904, pointing to regional variation among groups historically labeled as “Punic”. These data support a model in which male individuals at ADF descended from diverse but interconnected populations spanning the Mediterranean, Balkans, and western Asia, with gene flow likely occurring during the late Republic and Imperial periods (Figure 5).

In sum, the ancestry of ADF_m individuals appears shaped by multi-layered, male-driven recent demographic processes linked to Mediterranean, Balkan, and Near Eastern groups, likely first introduced during the Iron Age and especially the Roman Imperial period, revealing how this community was genetically transformed by the broader dynamics of Imperial mobility and demographic integration.

### Diverging worlds in the same necropolis

Mitochondrial and Y-chromosome haplogroups were successfully assigned for the majority of individuals with sufficient coverage (Supplementary Table 1). Among ADF_f individuals, mitochondrial haplogroups include lineages within U and H, specifically U5a, U3b, U1, K2a, H34, and H16. These haplogroups are broadly distributed across Eurasia, with notable associations to Eastern Europe, the Caucasus, and the Near East. In contrast, ADF_m individuals predominantly carry mitochondrial haplotypes belonging to subclades of haplogroup H, which is widespread across Europe and the Mediterranean. Y-chromosome haplogroups were successfully determined for three ADF_m individuals. These include J-BY37605, a subclade of haplogroup J-M304, as well as R-Y52 and E-CTS1273. These lineages are broadly distributed across the Near East, Mediterranean, and Eurasian Steppe, indicating diverse paternal ancestries within the ADF sample. Overall, the uniparental markers support the patterns observed in genome-wide analyses, indicating heterogeneous maternal and paternal lineages within the ADF population.

Taken together, the genetic results depict two partially distinct ancestries coexisting in the same population: ADF_f individuals show stronger genetic affinities to Eastern Europe, the Steppe, and the Armenian Highlands, suggesting their genetic ancestry is linked to mobile pastoralist or trans-Eurasian groups. Conversely, ADF_m individuals align more with Punic, Roman, and Mediterranean sources, thus linked to broader patterns of imperial mobility.

The contrasting ancestries of ADF_m and ADF_f individuals reflect the intersection of continental and imperial demographic forces acting across Eurasia from the Late Bronze Age through the Roman period. While Iron Age societies of the Balkans and the Steppe were often characterized by regional endogamy and genetic continuity, the expansion of the Roman Empire introduced new axes of mobility, fostering long-distance migration, military resettlement, and colonial infrastructure that reshaped the genetic landscapes of inland regions far from the Mediterranean coast. In this context, ADF functioned as a continental contact zone, absorbing gene flow from both eastern Steppe-related populations and western and southern groups connected to Roman and Punic spheres of influence. Together, these sex-specific patterns illustrate how different forms of mobility intersected in provincial communities. ADF thus emerges not as a peripheral outpost but as a genetic and cultural crossroad, where diverse ancestries were integrated through asymmetric social and political processes tied to both regional traditions and the expanding reach of Roman power.

## Discussion

The genetic analysis of the ADF population from Roman Dacia provides insight into the demographic processes and cultural transformations that unfolded during the Roman conquest and subsequent colonization of the region. Our findings reveal a complex interplay of local Dacians, Roman settlers, and neighboring nomadic groups, which collectively shaped the genetic landscape of this frontier province. This is consistent with the broader nature of the Roman Empire, a vast and diverse entity, stretching from Britain in the northwest to Egypt in the southeast. At its height, the Empire encompassed a multitude of cultures, languages, and ethnicities, all unified under Roman rule^7,8^.

The integration of these diverse communities has often been considered under the heading of ‘Romanization’. However, recent scholarship has moved away from interpreting the annexation of territories by the Roman Empire as a uniform process of assimilation into a homogeneous Roman culture, instead emphasizing the diversity of identities across the Roman provinces^9–13^. Local communities often retained aspects of their cultural practices or developed hybrid or “creolized”^14^ identities under Roman rule. Such interactions were not one-sided, however, as Romans also adopted elements from local populations in a reciprocal process of cultural exchange. For reasons of convenience, the present study nevertheless occasionally retains the term of Romanization as a shorthand term for this complex reconfiguration, while acknowledging that the debate over the term remains unresolved, that it has different resonances in different academic traditions, and that no fully satisfactory alternative has yet emerged^15^.

The Roman legions were the backbone of the Empire’s expansion and defense, and their presence in the provinces contributed significantly to the cultural, political, social and linguistic reconfiguration of conquered populations^16^. In frontier regions such as Roman Dacia, which was heavily militarized following the massive deployment of military force in its conquest, soldiers not only defended the borders but also contributed to the spread of Roman culture, law and infrastructure. Moreover, after completing their service, many soldiers were granted land in the provinces as part of their veteran settlements. These colonies, such as Ulpia Traiana Sarmizegetusa, became centers of Roman administration and culture, with Roman law, architecture, and urban planning being introduced to the local population^17^. In parallel, the military camps, such as those in Apulum, often attracted civilian settlements known as *canabae*. These settlements housed traders, craftsmen, and families of soldiers, creating a symbiotic relationship between the military and civilian populations. Over time, these *canabae* grew into thriving towns, further spreading Roman culture and economic practices^18^.

This complex social landscape shaped by soldiers, settlers, merchants, and local people provides an important context for interpreting the genomic findings from ADF. Our results reveal contrasting male and female genetic ancestries, distinct patterns of genetic affinity, and the enduring impact of both Roman- and Steppe-related lineages. The ADF_f individuals (females) exhibit strong genetic affinity with Steppe-related and Central Asian Iron Age populations. This pattern is consistent either with the integration of females from these groups into local Dacian communities or with the persistence of local female lineages that already carried genetic affinities to Steppe-Central Asian populations prior to the Roman conquest. The dominance of Component 2 in ADF_f individuals, which is strongly represented in Steppe and Central Asian populations, indicates a substantial contribution of Steppe-related ancestry to the female genetic profile in ADF.

Within the complex demographic landscape of the Apulum area, local women could have played a particularly significant role in shaping the reconfigured communities of the province, often serving as a bridge between Roman and local cultures; a similar model has been previously advanced for other culturally heterogeneous groups, such as the Etruscans^19^. Local women in the provinces were central to the social and cultural fabric of their communities^20^. Women’s experiences were likely diverse, including enslavement and violence^21^ as a result of the conquest, but even though their roles varied depending on their social status, ethnic background, and regional context, they often acted as important agents in mediating between Roman and local cultures, particularly through marriage, family life and religious practices^22^.

Roman soldiers were stationed far from their homes and often formed relationships with local women, leading to the creation of mixed families^23^. These mixed families likely contributed to the cultural assimilation of the provinces, as they often adopted Roman customs while retaining elements of their local heritage^24^. Indeed, the Roman people in the provinces also contributed to religious syncretism, with Roman gods being worshipped alongside local deities^25^. Moreover, local women were also active participants in religious life, both in Roman and local cults; for instance, the worship of Zalmoxis, a Dacian deity related with some practices by Siberian peoples^26^, continued alongside the Roman pantheon in the province, with women playing a key role in maintaining these religious traditions^27^. The presence of local women and their public role in religious activity within the province is also supported by epigraphy^28,29^. Inscriptions often mention local women, particularly in the context of family relationships and religious dedications^30,31^. These inscriptions provide evidence of the social integration of some local women into Roman society, as well as their continued engagement local traditions, but this is limited to the wealthier minority able to afford inscriptions and to participate in Roman-style dedications, using Latin. The ancient genomic data now provide evidence for the presence and continuity of a broader segment of the female population, extending beyond those visible in the epigraphic record.

Our data show that the ADF_f individuals show a strong genetic connection to populations from the Caucasus, particularly Armenia, which represented the last great effort of Rome to bring the Pontic area under control. This suggests that the Caucasus region could have served as a genetic corridor linking Eastern Europe and Western Asia, facilitating the movement of peoples and genes into Roman Dacia^32^. The high outgroup f3-statistics values between ADF_f and Caucasus populations indicate a deep ancestral connection, possibly reflecting gene flow or shared ancestry with Armenian Highland groups during the Late Bronze Age and Iron Age. Similarly, the ADF_f individuals show significant genetic affinities with Eastern European and Steppe populations, such as *Russia_LBA*, *Moldova_LBA*, and *Tajikistan_Ksi*. This supports the idea that Steppe-related ancestry was a major component of the Dacian gene pool, likely introduced through interactions with Scythian or Sarmatian groups before its conquest by the Romans^33^. The Sarmatians, in particular, were known to have inhabited the Pontic-Caspian Steppe and Carpathian regions^34^, and their genetic influence on Roman Dacia is evident in the ADF_f individuals’ genetic profile.

In contrast, the ADF_m (male) individuals show a more heterogeneous genetic profile, with significant contributions from Mediterranean and North African populations, as indicated by the elevated Component 1 and Component 3 in the admixture analysis. This suggests that male-mediated gene flow from Roman settlers and Mediterranean populations was a key factor shaping the genetic makeup of Roman Dacia. The presence of Component 3, which is rare in Steppe and Central Asian groups, but common in Punic- and Phoenician-associated populations, further supports the idea that male ancestors of the ADF population may have originated from Mediterranean sources. This is consistent with historical accounts of Roman colonization of the area, which often involved the settlement of Roman soldiers and veterans from diverse regions of the Roman Empire in newly conquered territories^35^. Historical sources such as Flavius Eutropius (*Breviarium ab Urbe condita* VIII.6.2) describe Dacia as populated by large numbers of people transferred “from across the whole Roman world”^7^. These people shaped the economic, military, and cultural life of the provinces, contributing to the Romanization process while also introducing new cultural elements from their homelands^3^. Roman settlers and veterans were among the most prominent allochthonous groups in the provinces. After completing their military service, many Roman soldiers were granted land in the provinces, where they established veteran colonies and agricultural estates^36^. These settlers brought with them Roman agricultural practices, legal systems, and cultural traditions, which were gradually adopted by the local population.

However, mobility into Roman Dacia was not limited to military personnel. Merchants, traders, specialized workers, enslaved individuals and other non-military individuals played a central role in the economic and cultural integration of the province, facilitating the circulation of goods, ideas, and people across the Empire^37^. The prominence of settlements such as Alburnus Maior (modern Roșia Montană), a major gold-mining center in the Apuseni Mountains, reflects Dacia’s incorporation into extensive imperial trade networks^38–41^. Its prosperity depended not only on mineral resources but also on well-developed infrastructure, including road systems connecting the region to key military, administrative, and commercial hubs across the Danubian frontier, the Balkans, and the wider Mediterranean^42^. These networks enabled the efficient movement of resources, officials, and commodities, while fostering urban development and cultural exchange^43,44^. Epigraphic evidence from Alburnus Maior, including inscriptions referencing individuals from Italy, the Balkans, and eastern provinces, further attests to the region’s demographic diversity^45^. In addition, enslaved individuals and freedmen, often involved in mining, agriculture, and infrastructure development, contributed significantly to the economic and social fabric of the province^46–49^. Together, these observations suggest that Roman Dacia received an influx of people from different regions. In this context, the ADF_m genomic profile likely reflects male-biased contributions from multiple sources and exemplifies the demographic impact of Roman colonization on the region. This is further supported by D-statistics, which indicate that ADF_m shares more recent or extensive genetic drift with Mediterranean populations than with ADF_f, highlighting a sex-biased pattern of genetic affinities associated with the broader demographic footprint of Roman colonization.

We note that the observed sex-biased patterns might be influenced by sampling bias or cemetery structure. However, the genomic patterns we identified in that peripheral Roman population align with historical accounts of Roman colonization, which frequently involved the settlement of Roman soldiers and veterans in newly conquered territories alongside the integration of local women into the localsociety^50^.

The genomic patterns also enrich the understanding of population change. The genetic diversity of the ADF sample also reflects the cultural and genetic complexity of Roman Dacia, which served as a frontier region at the intersection of Roman, Dacian, and nomadic influences. The Roman conquest and subsequent colonization of Dacia likely facilitated the integration of diverse genetic components into the local population, resulting in a heterogeneous genetic landscape that persisted even after the Roman withdrawal in 271 CE^51^.

The distribution of uniparental markers is consistent with the genome-wide results, supporting the presence of diverse maternal and paternal lineages within the ADF sample. Mitochondrial haplogroups observed in ADF_f individuals are broadly associated with Eastern Europe, the Caucasus, and the Near East^52–57^, whereas ADF_m individuals predominantly carry lineages widely distributed across Europe and the Mediterranean^58^. In addition, Y-chromosome haplogroups point to heterogeneous paternal origins spanning the Near East, the Eurasian Steppe, and North Africa^59–61^. Together, these patterns further support multiple, geographically diverse genetic contributions shaping this community.

Our findings provide biological insight into the broader implications of Roman imperial policy and its long-term demographic and cultural impacts on frontier provinces. The Roman Empire’s strategy of colonization and integration extended beyond the establishment of military garrisons and administrative structures: it also involved the intentional movement and settlement of diverse populations as a means of consolidating control. In the case of Roman Dacia, the evidence of Mediterranean and North African genetic affinities foregrounds how male-biased contributions may have played an important role in the Romanization process. This likely reflects the settlement of Roman veterans, officials, merchants, and their families, who contributed not only to the administrative and economic development of the province but also to the region’s genetic makeup. The resulting demographic heterogeneity illustrates how the Empire’s expansion was accompanied by complex patterns of human mobility and cultural integration. Importantly, this was not an isolated phenomenon: similar patterns have been documented in other Roman provinces such as Britain and Gaul, where military colonies and veteran settlements served as vectors of Roman cultural linguistic^62–64^ and genetic influence. These observations underscore the Empire’s capacity directly and indirectly to reshape both the cultural and, as argued here, the genetic makeup of its provinces through both structural and biological means, offering valuable insights into the mechanisms of Imperial cohesion and legacy.

However, the integration of local populations into the Romanized society was not uniform and often involved sex-biased patterns of population genetic affinity. The presence of local women and allochthonous people in Roman Dacia was a key factor in shaping the cultural landscape of Roman Dacia. Local women likely contributed to social integration, while also maintaining local traditions in religion and craftsmanship. Allochthonous people, including Roman settlers, merchants, and enslaved individuals, contributed to the economic and cultural life of the province, introducing new practices and traditions from across the Roman Empire. Together, these groups facilitated a cultural exchange that created a dynamic and diverse social landscape in the Roman provinces.

Similar patterns of increased genetic heterogeneity and long-distance mobility have been reported, albeit to varying degrees, in other Roman communities, including those in Britain^65–68^ and Italy^8,69–71^, where genomic data reveal individuals with diverse geographic origins. In this context, the patterns observed at ADF are broadly consistent with the dynamics associated with Roman expansion, while also reflecting the specific historical and geographic setting of the Dacian frontier.

Our findings, consistent with previous studies, suggest that Roman colonization was a multifaceted process involving both demographic and cultural integration. However, because they are based on genomic data, these findings are independent of the inscriptions on which previous characterizations have relied, which were limited to a minority of the population. Local women appear to have played a central role in maintaining the genetic continuity of frontier populations – at least in the case of Dacia – while incoming Roman settlers may have contributed disproportionately to the male genetic pool. Such sex-biased patterns of genetic affinity likely reflect the social dynamics of Roman expansion, including intermarriage, assimilation, and the establishment of settler communities following the demographic impact of violent conquest.

Despite the limited sample size, our study provides one of the first genome-wide perspectives on population dynamics in Roman Dacia, revealing complex and sex-biased admixture patterns. These patterns highlight differential genetic contributions from Steppe-related, Caucasus, and Mediterranean-associated populations, reflecting the interplay of Roman colonization, local Dacian communities, and neighboring nomadic groups. We acknowledge that the observed sex-biased signals may be influenced by sampling bias or necropolis-specific structure, and therefore should be interpreted with caution. In particular, a single necropolis, particularly in a context such as Apulum that likely included a high proportion of incomers, may not be representative of Roman Dacia as a whole. Nevertheless, the ADF dataset offers a preliminary framework for exploring the demographic history of this community, complementing but extending beyond models from inscriptions and historical sources. While the identified patterns are consistent with historical models of Roman colonization involving mobility and integration across the Empire, additional data from other sites will be necessary to assess their wider applicability.

Overall, our findings have broader implications for understanding the impact of Roman imperial expansion on frontier communities, highlighting the complex social dynamics of colonization and the integration of local populations within the Empire. While our results are limited to a single site, they underscore the potential of genomic data to illuminate patterns of mobility and interaction in Roman frontier regions. Future studies incorporating larger datasets from multiple sites will be essential to further resolve the genetic diversity of these populations and to better characterize the demographic processes that shaped the Roman Empire and its legacy in Europe and beyond.

## Methods

### Archeological site

Roman Dacia has yielded data from the systematic or partial investigation of approximately 50 necropolises and funerary sites, though the extent and depth of information vary widely^35,72^. Archaeological research identified two large bi-ritual necropolises, with inhumation and cremation burials coexisting without topographical separation in Apulum, present-day Alba Iulia, Romania (Figure 6)^73–75^. They have been used throughout the 2^nd^ and 3^rd^ centuries CE: the northern necropolis in the Stadium area and the southern necropolis on *Dealul Furcilor* (also known as Forks Hill or Podei)^76,77^. These necropolises align with the Roman custom of placing burial grounds along roads outside urban centers. Among these, the Alba Iulia-*Dealul Furcilor* (ADF) necropolis, spanning 35 hectares, stands out as the largest known funerary complex in Dacia, with only 20-25% excavated by the year 2011, and 1,243 in-situ burials recovered by 2018, yet it remains incompletely explored^72^.

**Figure 6:**
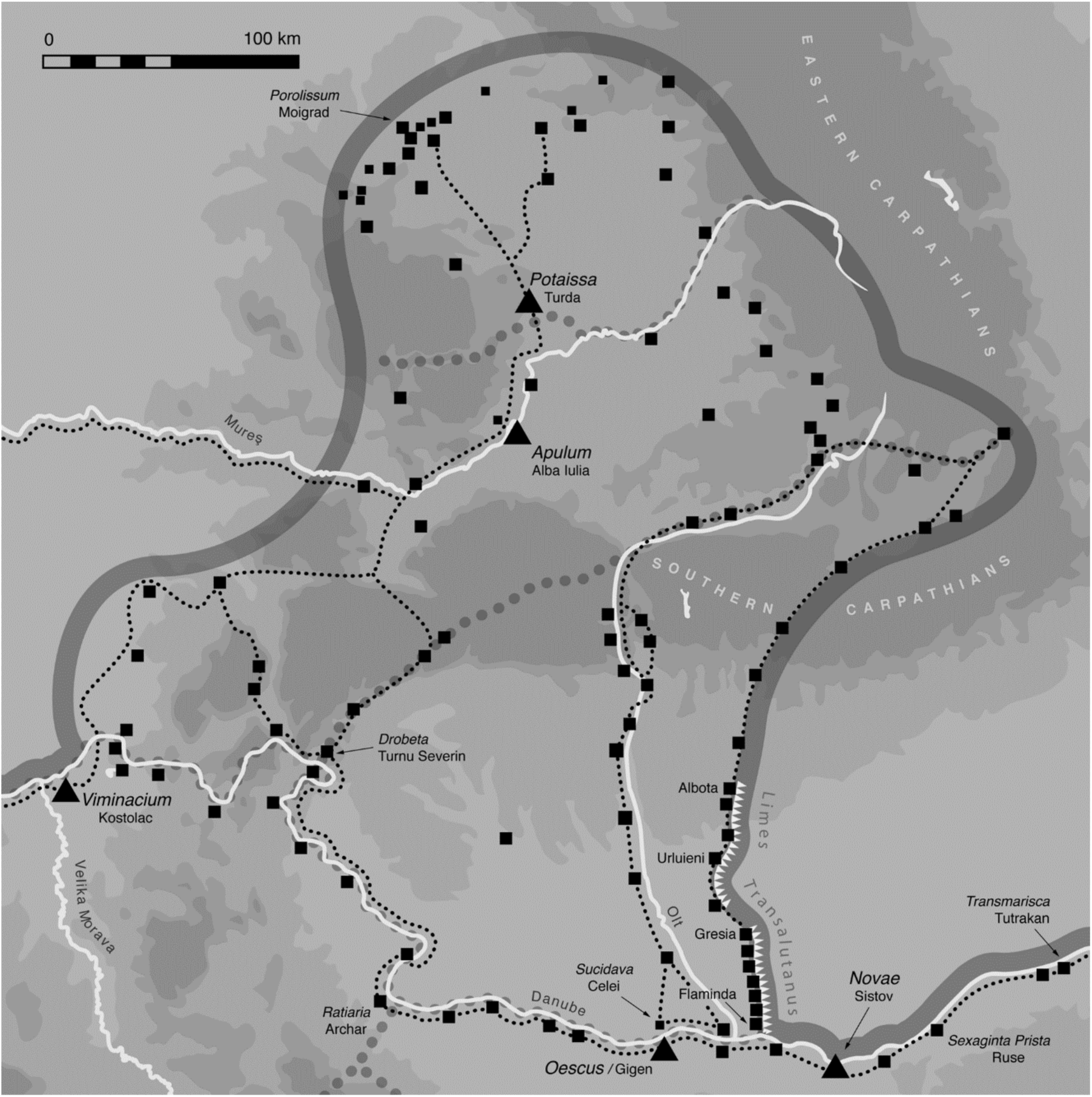
Location of Apulum in the context of Roman Dacia. From *Danube Limes – UNESCO World Heritage* (Pen & Sword; CHC, University of Salzburg), by David Breeze and Kurt Schaller. Reproduced with permission of ao. Univ.-Prof. i.R. Dr. Andreas Schwarcz.

Topographically, Apulum represents the most extensive and densely populated urban conurbation of Roman Dacia. It comprised two distinct yet coexisting urban centers situated approximately 2 km apart^78,79^. The first, Apulum I, developed from the *pagus* founded during Trajan’s reign and originally subordinate to *Colonia Ulpia Traiana Sarmizegetusa*. Over time, it evolved into the *Municipium Aurelium Apulense* under Marcus Aurelius and subsequently attained colonial status as *Colonia Aurelia Apulensis*. Its prosperity was such that, by the mid-third century CE, under Emperor Trebonianus Gallus, it bore the honorary title “*Chrysopolis*” – the “City of Gold”^80^.

The second urban nucleus, Apulum II, originated from the *canabae* located southwest of the castrum of Legio XIII Gemina and was granted the status of *Municipium Septimium Apulense* under Septimius Severus^78^. While Apulum II’s development was closely tied to its military function and its social composition reflected a limited number of *equites* and *collegia*, Apulum I displayed evidence of numerous villas and professional associations, suggesting a market-oriented economy. Notably, a higher proportion of *homines novi* – new men of elevated social mobility – appears attested at Apulum II compared to Apulum I, highlighting distinct socio-economic dynamics between the two centers^81^.

The ADF necropolis likely originated as a burial ground for *Municipium Aurelia Apulensis* – the former administrative centre established under Marcus Aurelius – with graves located along the road connecting the municipium to the Roman fort and *canabae*. To date, approximately 1,200 bi-ritual graves have been identified. Thirty-four skeletal individuals were sampled for genetic analysis in this study. Among these, 44,1% were classified as adult males and 38,2% as adult females, with the remaining individuals being juveniles whose biological sex could not be determined solely based on osteological traits, due to the absence of developed dimorphism. The selection of samples for this study prioritized the visual preservation of the osteological remains, with classification further informed by deposition patterns, taphonomic factors, and associated grave goods, including coins, amulets, *unguentaria*, and nails from *caligae* (Roman shoes), lamps, jewellery. Although only three individuals have been directly radiocarbon dated, the results support a chronology centred on the 2^nd^-3^rd^ centuries CE. Individual 21 (ADF_04; Poz-145332; 1880 ± 30 BP) yielded a calibrated date of 110–236 CE (90.3% probability), individual 22 (ADF_1; Poz-118945; 1830 ± 30 BP) a date of 117–252 CE (91.5% probability), and individual 30 (ADF_2; Poz-118946; 1760 ± 30 BP) a date of 211–383 CE (93.1% probability). The presence of coins in the burials further supports this chronology, whereas the remaining individuals were dated on the basis of their archaeological context.

### Ancient DNA extraction, library preparation, and sequencing

We screened total of 34 teeth from the ADF necropolis for the presence of aDNA. Laboratory work was conducted in dedicated clean-room facilities at the Paleogenomics Laboratory, University of California, Santa Cruz (UCSC-PGL), under stringent contamination-prevention measures. Teeth were mechanically brushed to remove soil and treated with a 3% Sodium Hypochlorite solution to clean the outer surfaces. After rinsing in molecular biology-grade water and 95% ethanol, the teeth were dried and exposed to UV radiation for 10 minutes on each side to minimize surface contamination. DNA was extracted from the cellular cementum of tooth roots using a minimally destructive protocol for DNA extraction^82^. DNA extracts were used to construct paired-end, double-indexed single-stranded genomic DNA (ssDNA) sequencing libraries^83^. Libraries underwent half-UDG treatment to partially remove deaminated bases from aDNA fragments, balancing authenticity with data preservation. Sequencing was performed on Illumina NovaSeq 6000 platforms, generating 2×150 bp paired-end reads.

### Sequencing reads alignment

AdapterRemoval^84^ was used to trim adapters and collapse forward and reverse reads into single sequences, with the following parameters: `--minlength 30 --minquality 25 --trimns --trimqualities`. Reads were aligned to the human reference genome hg19 using BWA^85^ (version 0.7.17) with parameters `mem -k19 -r 2.5`. The resulting SAM files were converted to BAM format using SAMtools^86^. Reads with a mapping quality below 30 were filtered out, and duplicate reads were removed using DeDup^87^, retaining the highest-quality read.

### Ancient DNA authentication and genotyping procedures

The authenticity of aDNA was verified by the presence of an excess of C-to-T substitutions at the ends of sequencing reads, assessed using mapDamage^88^. To control for post-mortem damage, the three terminal bases on both sides of the reads were trimmed. Mitochondrial contamination was assessed using Schmutzi^89^, aligning reads to mitochondrial genomes of present-day individuals from around the world.

Biological sex was determined by comparing the number of reads mapped to the Y-chromosome with those mapped to autosomes or the X-chromosome^90^, using high-quality reads (mapping quality ≥ 30). X-chromosome contamination in male individuals was assessed using ANGSD^91^ (Supplementary Table 1).

To contextualize the ADF sample set within broader ancient and present-day populations, we performed a genomic principal component analysis (PCA) using ancient and modern Eurasian and North African samples from the AADR Human Origins (HO) dataset reported in AADR^92^. Furthermore, filters of a minimum base and mapping quality 30 (-min-BQ and -min-MQ options) were used via samtools mpileup command. Pseudo-haploid genotypes were called by pileupCaller (sequenceTools version 1.2.2: https://hackage.haskell.org/package/sequenceTools) picking a random read at each site^93^.

Mitochondrial consensus sequences were obtained from reads with mapping quality ≥ 30, and haplogroups were assigned using Haplogrep2^94^ (Phylotree Build 17). Y-haplogroups for male individuals were identified using Yleaf v3.1^95^.

### Biological relatedness inference

The degree of biological relatedness among the sampled individuals from ADF was estimated using TKGWV2^96^, a pipeline developed for ultra-low coverage ancient DNA data. The method evaluates genome-wide allele sharing between pairs of individuals while incorporating external allele frequency information to estimate relatedness coefficients.

### Dataset compilation

Data pertaining HO SNP panel from published ancient (n = 888) and present-day (n = 2,267) individuals^8,60,69,70,92, 97–140^ were obtained from the AADR v54.1 repository^92^ and combined with 14 newly sampled ADF individuals sharing at least 10,000 variants with the panel. Present-day data included populations from Western Eurasia, Carpathian region, and Africa (Yoruba). Ancient samples focused on both Mediterranean and Caspian Sea regions.

### Population genetic analysis

Principal Component Analysis (PCA) was performed using the smartpca program from the EIGENSOFT^141^ package, projecting ancient samples into the principal component space of present-day populations (160,290 variants). The analysis used default settings with the “`lsqproject: YES`” option. Population structure was explored using ADMIXTURE^142^, with variants in moderate to high linkage disequilibrium (r² > 0.4), SNPs with >60% missing data, and minor allele frequencies <5% removed using PLINK^143^. Unsupervised ADMIXTURE analysis was performed for K values ranging from 2 to 10. Genetic similarity between populations was assessed using outgroup-f3 statistics calculated with qp3Pop (AdmixTools^144^). D-statistics were used to explore genetic relationships among four populations, calculated in the form of D(ADF, X; Y, Outgroup).

## Data availability

The sequencing data generated in this study have been deposited in the European Nucleotide Archive (ENA) under accession number [to be added upon acceptance]. Previously published genomic data used in this study are available through the Allen Ancient DNA Resource (AADR) repository^92^.

## Acknowledgements

The authors gratefully acknowledge ao. Univ.-Prof. i.R. Dr. Andreas Schwarcz (Institut für Österreichische Geschichtsforschung, Universität Wien, Austria) for kindly granting permission to access and reproduce maps from the Danube Limes - UNESCO World Heritage Project.

## Authors contributions

F.D.A.: Conceptualization, Methodology, Formal analysis, Investigation, Data curation, Writing – original draft, Writing – review & editing. I.B.: Conceptualization, Investigation, Resources, Writing – review & editing. K.K.: Investigation, Methodology; A. C. B. – Investigation, Resources, Writing – review & editing. A.T.: Investigation, Resources, Writing – review & editing. J.P.: Investigation, Writing – review & editing. M.G.: Conceptualization, Investigation, Resources, Writing – review & editing. L.F.-S.: Supervision, Writing – review & editing. C.E.G.A.: Conceptualization, Supervision, Funding acquisition, Writing – review & editing. All authors reviewed and approved the final manuscript.

## Ethical approval

All necessary permits for sampling and genetic analysis were obtained from the Alba County Directorate for Culture. The export of samples for ancient DNA analysis was conducted in accordance with national regulations and approved by the relevant authorities. Laboratory and analytical procedures followed established ethical standards for research involving ancient human remains.

## Supplementary tables and figures legends

**Supplementary Table S1: Sequencing statistics, authentication metrics, and metadata for ADF genomic libraries.** Sequencing and mapping statistics for all ADF libraries screened for ancient DNA (aDNA). “Archeological_ID” refers to the archaeological sample identifier and “Lab_ID” to the laboratory code. The archeological ID should be used for references to the data presented in this study. Sequencing metrics include “Read count” (total sequencing reads), “Mapped reads” (mapped reads after duplicate removal), “Endogenous DNA (%)” (endogenous DNA content), “Mean DoC” (mean genome-wide depth of coverage), and “1240k_HO cov” (average coverage on the 1240K_HO SNP panel). Sex determination using the R_y method includes “Nseqs” (number of reads used for sex determination), “NchrX+NchrY” (reads mapping to both chromosomes X and Y), “NchrY” (reads mapping to chromosome Y), “R_y” (ratio of Y-chromosome reads), “SE” (standard error), and “95% CI” (95% confidence interval), followed by the resulting assignment. Information on the uniparental markers include “mt_coverage” (mitochondrial DNA coverage), “mt Hg” (mitochondrial haplogroup assignment), and “Y-chr Hg” (Y-chromosome haplogroup). “Quality” indicates haplogroup assignment quality as assessed by Haplogrep3, and “Range” indicates the mtDNA regions used for haplogroup determination and “Y Hg markers” the corresponding for the Y-chromosome. Post-mortem damage patterns are reported as “1 pos 5pC>T”, “2 pos 5pC>T”, and “3 pos 5pC>T”, representing the frequency of C→T substitutions at positions 1, 2, and 3 from the 5′ end of sequencing reads, respectively.

**Supplementary Table S2: Pairwise kinship relationships among ADF individuals.** Pairwise relatedness estimates among ADF individuals inferred using TKGWV2 (see Methods). Sample: First individual of tested pair; Sample2: Second individual of tested pair; Used_SNPs: Number of SNPs used to infer relatedness; HRC: Halved Relatedness Coefficient (Unrelated < 0.0625; 2nd Degree between 0.0625 and 0.1875; 1st Degree > 0.1875); counts0: Number of non-shared alleles; counts4: Number of shared alleles; Relationship: Descriptive relationship based on HRC value.

**Supplementary Figure S1:** Scree plot showing the cross-validation error across clustering solutions and supporting the choice of the optimal number of clusters in the ADMIXTURE analysis.

## References

1. Fulford, M. Territorial Expansion and the Roman Empire. World Archaeol. 23, 294–305 (1992).

2. Vekony, G. Dacians Romans Romanians. (Matthias Corvinus, New York, 1999).

3. Varga, R. THE NATURE OF ROMAN DOMINION OVER THE PROVINCE OF DACIA NOTES ON THE ROMANIZATION PHENOMENON AND ITS LIMITS. J. Anc. Hist. Archaeol. 8, (2021).

4. Pankova, S. & Simpson, S. J. Masters of the Steppe: The Impact of the Scythians and Later Nomad Societies of Eurasia : Proceedings of a Conference Held at the British Museum, 27-29 October 2017. (Archaeopress Archaeology, 2020).

5. Grumeza, I. *Dacia: Land of Transylvania, Cornerstone of Ancient Eastern Europe*. (Hamilton Books, Lanham, Md Plymouth, 2009).

6. Silva, M. et al. An individual with Sarmatian-related ancestry in Roman Britain. Curr. Biol. CB 34, 204–212.e6 (2024).

7. Ethnicity and Identity in the Roman Empire. in Rome: An Empire of Many Nations: New Perspectives on Ethnic Diversity and Cultural Identity (eds Price, J. J., Finkelberg, M. & Shahar, Y.) 15–84 (Cambridge University Press, Cambridge, 2021).

8. Antonio, M. L. et al. Ancient Rome: A genetic crossroads of Europe and the Mediterranean. Science 366, 708–714 (2019).

9. Millett, M. The Romanization of Britain: An Essay in Archaeological Interpretation. (Cambridge University Press, 1990).

10. Woolf, G. Romanization 2.0 and its alternatives. Archaeol. Dialogues 21, 45–50 (2014).

11. Versluys, M. J. Understanding objects in motion. An archaeological dialogue on Romanization. Archaeol. Dialogues 21, 1–20 (2014).

12. Terrenato, N. The deceptive archetype. Roman colonialism and post-colonial thought. in Ancient Colonizations. Analogy, Similarity, and Difference 59–72 (Duckworth, London, 2005).

13. Mattingly, D. J. Imperialism, Power, and Identity: Experiencing the Roman Empire. (Princeton University Press, 2011).

14. Webster, J. Creolizing the Roman Provinces. Am. J. Archaeol. 105, 209–225 (2001).

15. Versluys, M. J. Getting out of the comfort zone. Reply to responses. Archaeol. Dialogues 21, 50–64 (2014).

16. The Discourse on Romanization in the Age of Empires. in Decolonizing Roman Imperialism: The Study of Rome, Romanization, and the Postcolonial Lens (ed. Lambert, D. H.) 15–50 (Cambridge University Press, Cambridge, 2024). doi:10.1017/9781009491044.002.

17. Alicu, D. Figured Monuments from Ulpia Traiana Sarmizegetusa: 55. (British Archaeological Reports Ltd, Oxford, 1979).

18. Sorin, N. Canabae legionis in Dacia. Military presence and municipal evolution at Apulum and Potaissa. ALFRED VON DOMASZEWSKI Lat. Epigr. Roman Emp. Acts Colloq. Held Timișoara Dec. 14th–17th 2022 https://www.academia.edu/122574506/Canabae_legionis_in_Dacia_Military_presence_and_municipal_evolution_at_Apulum_and_Potaissa (2024).

19. Daveloose, A. The Etruscan Woman and “Romanization”: An Onomastic Case Study. Etruscan Ital. Stud. https://doi.org/10.1515/etst-2024-0018 (2025) doi:10.1515/etst-2024-0018.

20. Ivleva, T., De Bruin, J. & Driessen, M. Embracing the Provinces: Society and Material Culture of the Roman Frontier Regions. (Oxbow Books, 2018). doi:10.2307/j.ctv13nb8qs.

21. Phang, S. Intimate conquests: Roman soldiers’ slave women and freedwomen. *Anc*. World 35, 207–237 (2024).

22. Noreña, C. F. Romanization in the Middle of Nowhere: The Case of Segobriga. Fragm. Interdiscip. Approaches Study Anc. Mediev. Pasts 8, (2019).

23. Busch, A. W. & Greene, E. M. Mother Courage and Her Children: The Family and Social Life of the Garrisons Stationed in Rome. in Women and the Army in the Roman Empire (eds Greene, E. M. & Brice, L. L.) 110–150 (Cambridge University Press, Cambridge, 2024). doi:10.1017/9781107705982.005.

24. Roselaar, S. T. Processes of Cultural Change and Integration in the Roman World. in (Brill, 2015).

25. Webster, J. Necessary Comparisons: A Post-Colonial Approach to Religious Syncretism in the Roman Provinces. World Archaeol. 28, 324–338 (1997).

26. Drugaş, Ş. G. P. The Name of Zalmoxis and Its Signiflcance in the Dacian Language and Religion. https://doi.org/10.3406/hiper.2016.914 (2016) doi:10.3406/hiper.2016.914.

27. Byros, G. Reconstructing Identities in Roman Dacia: Evidence from Religion. https://www.academia.edu/7256904/Reconstructing_Identities_in_Roman_Dacia_Evidence_from_Religion (2014).

28. Szabó, C. Women and Roman religion in Dacia: the epigraphic evidence. Cercet. Arheol. 31.**1**, 99–122 (2024).

29. Murzea, D. & Kruschwitz, P. Women of Roman Dacia (Re-)Centred: Roman Verse Inscriptions between Macro-History and Micro-Narrative. Mihăilescu-Bîrliba R Ardevan R Varga F Matei-Popescu O Țentea Eds Stud. Epigr. Hist. Honor. Ioannis Pisonis Wiesb. https://www.academia.edu/123359944/Women_of_Roman_Dacia_Re_Centred_Roman_Verse_Inscriptions_between_Macro_History_and_Micro_Narrative (2024).

30. Greene, E. M. The Role of Women in the Religious Activities of Roman Military Communities. in Women and the Army in the Roman Empire (eds Greene, E. M. & Brice, L. L.) 240–269 (Cambridge University Press, Cambridge, 2024). doi:10.1017/9781107705982.009.

31. Timea, V. T. Varga, The religion of the Dacians in the Roman Empire. Nemeti Dana Eds Dacians Roman Emp. Prov. Constr. Cluj-Napoca https://www.academia.edu/44873647/T_Varga_The_religion_of_the_Dacians_in_the_Roman_Empire (2019).

32. Bekker-Nielsen, T. Rome and the Black Sea Region: Domination, Romanisation, Resistance. (Aarhus University Press, 2006). doi:10.2307/j.ctv62hh2z.

33. Lopez, L. Dacia: A Contested Empire. https://www.academia.edu/16961424/Dacia_A_Contested_Empire.

34. Oţa, L. L. DACIANS OR SARMATIANS? TAMGA SIGNS IN DACIA (1ST CENTURY BC – 1ST CENTURY AD). Anc. Thrace Myth Real. Proc. Thirteen. Int. Congr. Thracology Kazanlak *Sept. 3 - 7 2017 Vol 2* https://www.academia.edu/108701448/DACIANS_OR_SARMATIANS_TAMGA_SIGNS_IN_DACIA_1ST_CENTURY_BC_1ST_CENTURY_AD_ (2023).

35. Mihailescu-Birliba, L. & Asăndulesei, A. ROMAN ARMY AND SALT EXPLOITATION IN DACIA. J. Anc. Hist. Archaeol. 6, (2019).

36. Ferjančić, S. Sirmium and the veterans of the Roman army. in *Vol* 8 Studia epigraphica et militaria: In memoriam Miroslava Mirković (eds Horster, M., Pelcer-Vujačić, O. & Ferjančić, S.) 13–24 (De Gruyter, 2024).

37. Lee, H. J. Transformative Trade: The Impacts of Trade on the Periphery of the Roman Empire. (Macquarie University, 2023). doi:10.25949/21992459.v1.

38. Cauuet, B. Alburnus Maior. in The Encyclopedia of Ancient History (John Wiley & Sons, Ltd, 2012). doi:10.1002/9781444338386.wbeah16003.

39. Damian, P. Alburnus Maior I. (CIMEC, Bucureşti, 2003).

40. Simion, M., Vleja, D. & Apostol, V. Alburnus Maior II. (CIMEC, Bucarest, 2005).

41. Damian, P. Alburnus Maior III. (CIMEC, 2008).

42. Glodariu, I. Dacian Trade with the Hellenistic and Roman World. (BAR Publishing, 1976). doi:10.30861/9780904531404.

43. Gudea, I. I., Somesan, T. & Pop, C. C. THE ROMAN LIMES IN TRANSYLVANIA – A GEOGRAPHICAL-ARCHAEOLOGICAL AXIS STRUCTURE. Transylv. Rev. Adm. Sci. 66, (2025).

44. Fodorean, F. THE ORIGIN AND DEVELOPMENT OF THE MAIN ROAD INFRASTRUCTURE AND THE CITY NETWORK OF ROMAN DACIA. J. Anc. Hist. Archaeol. 6, (2019).

45. Beu-Dachin, E. About the Greeks and the Greek Language in the written sources from Alburnus Maior. Acta Musei Napoc. https://www.academia.edu/43154688/About_the_Greeks_and_the_Greek_Language_in_the_written_sources_from_Alburnus_Maior (2015).

46. Harris, W. V. Demography, Geography and the Sources of Roman Slaves. J. Roman Stud. 89, 62–75 (1999).

47. Luciani, F. PUBLIC SLAVES IN ROME: ‘PRIVILEGED’ OR NOT? Class. Q. 70, 368–384 (2020).

48. Odochiciuc, A. & Mihailescu-Bîrliba, L. Occupations of Private Slaves in Roman Dacia. 231–247 http://saa.uaic.ro/occupations-of-private-slaves-in-roman-dacia/ (2014).

49. Vnukov, S. Overseas Trade in the Black Sea Region and the Formation of the Pontic Market from the First Century bce to the Third Century ce. in The Northern Black Sea in Antiquity: Networks, Connectivity, and Cultural Interactions (ed. Kozlovskaya, V.) 100–138 (Cambridge University Press, Cambridge, 2017). doi:10.1017/9781139094702.007.

50. Petropoulos, E. K. Review of: Dacia. Landscape, Colonisation, Romanisation. Routledge Monographs in Classical Studies. Bryn Mawr Class. Rev. https://bmcr.brynmawr.edu/2008/2008.09.10/.

51. Cocoş, R. et al. Genetic affinities among the historical provinces of Romania and Central Europe as revealed by an mtDNA analysis. BMC Genet. 18, 20 (2017).

52. Brotherton, P. et al. Neolithic mitochondrial haplogroup H genomes and the genetic origins of Europeans. Nat. Commun. 4, 1764 (2013).

53. Dawod, P. G. A. et al. Mutational Analysis and mtDNA Haplogroup Characterization in Three Serbian Cases of Mitochondrial Encephalomyopathies and Literature Review. Diagnostics 11, 1969 (2021).

54. Kerber, R. A., O’Brien, E., Munger, R., Smith, K. R. & Cawthon, R. M. Mitochondrial genetics of exceptional longevity in multigeneration matrilineages. 361899 Preprint at 10.1101/361899 (2018).

55. Kristjansson, D., Bohlin, J., Nguyen, T. T., Jugessur, A. & Schurr, T. G. Evolution and dispersal of mitochondrial DNA haplogroup U5 in Northern Europe: insights from an unsupervised learning approach to phylogeography. BMC Genomics 23, 354 (2022).

56. Nasidze, I. et al. Mitochondrial DNA and Y-chromosome variation in the caucasus. Ann. Hum. Genet. 68, 205–221 (2004).

57. Sequeira, J. J., Vinuthalakshmi, K., Das, R., van Driem, G. & Mustak, M. S. The maternal U1 haplogroup in the Koraga tribe as a correlate of their North Dravidian linguistic affinity. Front. Genet. 14, 1303628 (2024).

58. Hernández, C. L. et al. The distribution of mitochondrial DNA haplogroup H in southern Iberia indicates ancient human genetic exchanges along the western edge of the Mediterranean. BMC Genet. 18, 46 (2017).

59. Ilumäe, A.-M. et al. Phylogenetic history of patrilineages rare in northern and eastern Europe from large-scale re-sequencing of human Y-chromosomes. Eur. J. Hum. Genet. 29, 1510–1519 (2021).

60. Jeong, C. et al. A Dynamic 6,000-Year Genetic History of Eurasia’s Eastern Steppe. Cell 183, 890–904.e29 (2020).

61. Manco, L. et al. The Eastern side of the Westernmost Europeans: Insights from subclades within Y-chromosome haplogroup J-M304. Am. J. Hum. Biol. Off. J. Hum. Biol. Counc. 30, (2018).

62. Mullen, A. Multilingualism and multiple identities: interdisciplinary methodologies. in Southern Gaul and the Mediterranean: Multilingualism and Multiple Identities in the Iron Age and Roman Periods 1–144 (Cambridge University Press, 2013).

63. Mann, J. C. Legionary Recruitment and Veteran Settlement During the Principate. (Routledge, Boca Raton, FL, 2018).

64. Kelly, C. Conquest. in The Roman Empire: A Very Short Introduction (ed. Kelly, C.) 0 (Oxford University Press, 2006). doi:10.1093/actrade/9780192803917.003.0002.

65. Scheib, C. L. et al. Low Genetic Impact of the Roman Occupation of Britain in Rural Communities. Mol. Biol. Evol. 41, msae168 (2024).

66. Walton, A. et al. Beachy Head Woman: clarifying her origins using a multiproxy anthropological and biomolecular approach. J. Archaeol. Sci. 185, 106445 (2026).

67. Martiniano, R. et al. Genomic signals of migration and continuity in Britain before the Anglo-Saxons. Nat. Commun. 7, 10326 (2016).

68. Redfern, R. C., Marshall, M., Eaton, K. & Poinar, H. N. ‘Written in Bone’: New Discoveries about the Lives and Burials of Four Roman Londoners. Britannia 48, 253–277 (2017).

69. Antonio, M. L. et al. Stable population structure in Europe since the Iron Age, despite high mobility. eLife 13, e79714 (2024).

70. De Angelis, F. et al. First Glimpse into the Genomic Characterization of People from the Imperial Roman Community of Casal Bertone (Rome, First–Third Centuries AD). Genes 13, 136 (2022).

71. De Angelis, F. et al. Ancient genomes from a rural site in Imperial Rome (1st-3rd cent. CE): a genetic junction in the Roman Empire. Ann. Hum. Biol. 48, 234–246 (2021).

72. Dalmon. THE FUNERARY ARCHAEOLOGY OF ALBA IULIA-DEALUL FURCILOR. Bul. Cercurilor Științifice Studențești 26, 5–18 (2020).

73. Bolog, A. D. Necropola Romana de La Apulum, Dealul Furcilor – “Podei” (Campaniile 2008-2012) . (Risoprint, Cluj Napoca, 2017).

74. Protase, D. Necropola Oraşului Apulum. vol. Apulum, XII (1974).

75. Gligor, M., Bogdan, D., Mazare, P., Lipot, S. & Balteş, G. Roman burials in Apulum’s necropola from Dealul Furcilor. Terra Sebus 2, 117–139 (2010).

76. Bounegru, G. V. The Northern Necropolis of Apulum. “Ambulance Station” 1981-1985. (Mega Publishing House, Cluj-Napoca, 2017).

77. Bounegru, G. Roman Cemeteries from Apulum. Demarcation and Chronology, Scripta Classica. (Mega Publishing House, Cluj-Napoca, 2011).

78. Ardevan, R. & zerbini, L. La Dacia romana - Rubbettino editore. (Rubettino, 2007).

79. Ota, R. De la canabele legiunii a XIII-a Gemina la Municipium Septimium Apulense/From canabae legionis to Municipium Septimium Apulense. Canabele Legiunii XIII- Gemina Munic. Septimium ApulenseFrom Canabae Legion. Munic. Septimium Apulense https://www.academia.edu/45180876/De_la_canabele_legiunii_a_XIII_a_Gemina_la_Municipium_Septimium_Apulense_From_canabae_legionis_to_Municipium_Septimium_Apulense (2012).

80. Oltean, I. A. Dacia: Landscape, Colonization and Romanization. (Routledge, London, 2007). doi:10.4324/9780203945834.

81. Moga, V. PREFECŢI AI CASTRULUI LEGIUNII XIII GEMINA LA APULUM. Apulum 13, 651–657 (1975).

82. Harney, É. et al. A minimally destructive protocol for DNA extraction from ancient teeth. Genome Res. 31, 472–483 (2021).

83. Kapp, J. D., Green, R. E. & Shapiro, B. A Fast and Efficient Single-stranded Genomic Library Preparation Method Optimized for Ancient DNA. J. Hered. 112, 241–249 (2021).

84. Lindgreen, S. AdapterRemoval: easy cleaning of next-generation sequencing reads. BMC Res. Notes 5, 337 (2012).

85. Li, H. & Durbin, R. Fast and accurate short read alignment with Burrows–Wheeler transform. Bioinformatics 25, 1754–1760 (2009).

86. Li, H. et al. The Sequence Alignment/Map format and SAMtools. Bioinformatics 25, 2078–2079 (2009).

87. Peltzer, A. et al. EAGER: efficient ancient genome reconstruction. Genome Biol. 17, 60 (2016).

88. Jónsson, H., Ginolhac, A., Schubert, M., Johnson, P. L. F. & Orlando, L. mapDamage2.0: fast approximate Bayesian estimates of ancient DNA damage parameters. Bioinformatics 29, 1682–1684 (2013).

89. Renaud, G., Slon, V., Duggan, A. T. & Kelso, J. Schmutzi: estimation of contamination and endogenous mitochondrial consensus calling for ancient DNA. Genome Biol. 16, 224 (2015).

90. Skoglund, P., Storå, J., Götherström, A. & Jakobsson, M. Accurate sex identification of ancient human remains using DNA shotgun sequencing. J. Archaeol. Sci. 40, 4477–4482 (2013).

91. Korneliussen, T. S., Albrechtsen, A. & Nielsen, R. ANGSD: Analysis of Next Generation Sequencing Data. BMC Bioinformatics 15, 356 (2014).

92. Mallick, S. & Reich, D. The Allen Ancient DNA Resource (AADR): A curated compendium of ancient human genomes. Harvard Dataverse 10.7910/DVN/FFIDCW (2024).

93. Günther, T. & Nettelblad, C. The presence and impact of reference bias on population genomic studies of prehistoric human populations. PLoS Genet. 15, e1008302 (2019).

94. Weissensteiner, H. et al. HaploGrep 2: mitochondrial haplogroup classification in the era of high-throughput sequencing. Nucleic Acids Res. 44, W58–63 (2016).

95. Ralf, A., Montiel González, D., Zhong, K. & Kayser, M. Yleaf: Software for Human Y-Chromosomal Haplogroup Inference from Next-Generation Sequencing Data. Mol. Biol. Evol. 35, 1291–1294 (2018).

96. Fernandes, D. M., Cheronet, O., Gelabert, P. & Pinhasi, R. TKGWV2: an ancient DNA relatedness pipeline for ultra-low coverage whole genome shotgun data. Sci. Rep. 11, 21262 (2021).

97. Lazaridis, I. et al. The genetic history of the Southern Arc: A bridge between West Asia and Europe. Science 377, eabm4247 (2022).

98. Patterson, N. et al. Large-scale migration into Britain during the Middle to Late Bronze Age. Nature 601, 588–594 (2022).

99. Flegontov, P. et al. Palaeo-Eskimo genetic ancestry and the peopling of Chukotka and North America. Nature 570, 236–240 (2019).

100. Allentoft, M. E. et al. Population genomics of Bronze Age Eurasia. Nature 522, 167–172 (2015).

101. Mathieson, I. et al. The genomic history of southeastern Europe. Nature 555, 197–203 (2018).

102. Damgaard, P. de B., et al. 137 ancient human genomes from across the Eurasian steppes. Nature 557, 369–374 (2018).

103. Saag, L. et al. The Arrival of Siberian Ancestry Connecting the Eastern Baltic to Uralic Speakers further East. Curr. Biol. 29, 1701–1711.e16 (2019).

104. Gnecchi-Ruscone, G. A. et al. Ancient genomes reveal origin and rapid trans-Eurasian migration of 7th century Avar elites. Cell 185, 1402–1413.e21 (2022).

105. Gamba, C. et al. Genome flux and stasis in a five millennium transect of European prehistory. Nat. Commun. 5, 5257 (2014).

106. Mathieson, I. et al. Genome-wide patterns of selection in 230 ancient Eurasians. Nature 528, 499–503 (2015).

107. Zalloua, P. et al. Ancient DNA of Phoenician remains indicates discontinuity in the settlement history of Ibiza. Sci. Rep. 8, 17567 (2018).

108. Posth, C. et al. The origin and legacy of the Etruscans through a 2000-year archeogenomic time transect. Sci. Adv. 7, eabi7673 (2021).

109. Fernandes, D. M. et al. The spread of steppe and Iranian-related ancestry in the islands of the western Mediterranean. *Nat*. Ecol. Evol. 4, 334–345 (2020).

110. Marcus, J. H. et al. Genetic history from the Middle Neolithic to present on the Mediterranean island of Sardinia. Nat. Commun. 11, 939 (2020).

111. Gnecchi-Ruscone, G. A. et al. Ancient genomic time transect from the Central Asian Steppe unravels the history of the Scythians. Sci. Adv. https://doi.org/10.1126/sciadv.abe4414 (2021) doi:10.1126/sciadv.abe4414.

112. Narasimhan, V. M. et al. The Formation of Human Populations in South and Central Asia. Science 365, eaat7487 (2019).

113. Unterländer, M. et al. Ancestry and demography and descendants of Iron Age nomads of the Eurasian Steppe. Nat. Commun. 8, 14615 (2017).

114. Krzewińska, M. et al. Ancient genomes suggest the eastern Pontic-Caspian steppe as the source of western Iron Age nomads. Sci. Adv. https://doi.org/10.1126/sciadv.aat4457(2018) doi:10.1126/sciadv.aat4457.

115. Jeong, C. et al. Long-term genetic stability and a high-altitude East Asian origin for the peoples of the high valleys of the Himalayan arc. Proc. Natl. Acad. Sci. 113, 7485–7490 (2016).

116. Kılınç, G. M. et al. Human population dynamics and Yersinia pestis in ancient northeast Asia. Sci. Adv. https://doi.org/10.1126/sciadv.abc4587 (2021) doi:10.1126/sciadv.abc4587.

117. Veeramah, K. R. et al. Population genomic analysis of elongated skulls reveals extensive female-biased immigration in Early Medieval Bavaria. Proc. Natl. Acad. Sci. U. S. A. 115, 3494–3499 (2018).

118. Sikora, M. et al. The population history of northeastern Siberia since the Pleistocene. Nature 570, 182–188 (2019).

119. Mathieson, I. et al. Genome-wide patterns of selection in 230 ancient Eurasians. Nature 528, 499–503 (2015).

120. Wang, C.-C. et al. Genomic insights into the formation of human populations in East Asia. Nature 591, 413–419 (2021).

121. Olalde, I. et al. The genomic history of the Iberian Peninsula over the past 8000 years. Science https://doi.org/10.1126/science.aav4040 (2019) doi:10.1126/science.aav4040.

122. Liu, C.-C. et al. Ancient genomes from the Himalayas illuminate the genetic history of Tibetans and their Tibeto-Burman speaking neighbors. Nat. Commun. 13, 1203 (2022).

123. Aneli, S. et al. The Genetic Origin of Daunians and the Pan-Mediterranean Southern Italian Iron Age Context. Mol. Biol. Evol. 39, msac014 (2022).

124. Scorrano, G. et al. Bioarchaeological and palaeogenomic portrait of two Pompeians that died during the eruption of Vesuvius in 79 AD. Sci. Rep. 12, 6468 (2022).

125. Gelabert, P. et al. Genomes from Verteba cave suggest diversity within the Trypillians in Ukraine. Sci. Rep. 12, 7242 (2022).

126. Moots, H. M. et al. A genetic history of continuity and mobility in the Iron Age central Mediterranean. *Nat*. Ecol. Evol. 7, 1515–1524 (2023).

127. Jeong, C. et al. The genetic history of admixture across inner Eurasia. *Nat*. Ecol. Evol. 3, 966–976 (2019).

128. Lazaridis, I. et al. Ancient human genomes suggest three ancestral populations for present-day Europeans. Nature 513, 409–413 (2014).

129. Patterson, N. et al. Ancient admixture in human history. Genetics 192, 1065–1093 (2012).

130. Lazaridis, I. et al. Genomic insights into the origin of farming in the ancient Near East. Nature 536, 419–424 (2016).

131. Biagini, S. A. et al. People from Ibiza: an unexpected isolate in the Western Mediterranean. Eur. J. Hum. Genet. EJHG 27, 941–951 (2019).

132. Bergström, A. et al. Insights into human genetic variation and population history from 929 diverse genomes. Science 367, eaay5012 (2020).

133. Damgaard, P. de B., et al. 137 ancient human genomes from across the Eurasian steppes. Nature 557, 369–374 (2018).

134. Raghavan, M. et al. Upper Palaeolithic Siberian genome reveals dual ancestry of Native Americans. Nature 505, 87–91 (2014).

135. Raghavan, M. et al. Genomic evidence for the Pleistocene and recent population history of Native Americans. Science 349, aab3884 (2015).

136. Meyer, M. et al. A High-Coverage Genome Sequence from an Archaic Denisovan Individual. Science https://doi.org/10.1126/science.1224344 (2012) doi:10.1126/science.1224344.

137. Auton, A. et al. A global reference for human genetic variation. Nature 526, 68–74 (2015).

138. Prüfer, K. et al. The complete genome sequence of a Neanderthal from the Altai Mountains. Nature 505, 43–49 (2014).

139. Järve, M. et al. Shifts in the Genetic Landscape of the Western Eurasian Steppe Associated with the Beginning and End of the Scythian Dominance. Curr. Biol. 29, 2430–2441.e10 (2019).

140. Harney, É. et al. A minimally destructive protocol for DNA extraction from ancient teeth. Genome Res. 31, 472–483 (2021).

141. Patterson, N., Price, A. L. & Reich, D. Population Structure and Eigenanalysis. PLOS Genet. 2, e190 (2006).

142. Alexander, D. H. & Lange, K. Enhancements to the ADMIXTURE algorithm for individual ancestry estimation. BMC Bioinformatics 12, 246 (2011).

143. Purcell, S. et al. PLINK: A Tool Set for Whole-Genome Association and Population-Based Linkage Analyses. Am. J. Hum. Genet. 81, 559–575 (2007).

144. Petr, M., Vernot, B. & Kelso, J. admixr—R package for reproducible analyses using ADMIXTOOLS. Bioinformatics 35, 3194–3195 (2019).

