## Supplemental Note S1 for "On the Edge of Empire: Paleogenomic Insights into Roman Dacia"

### Supplementary Note S1

#### Historical Background

Conquered by Emperor Trajan in 106 CE after two major campaigns in 101—102 and 105—106 CE, Dacia represents the last territory to have been lastingly incorporated into the Roman Empire before its loss under Emperor Aurelian around 271 CE.[1] The establishment of Dacia as a Roman province marked a pivotal moment in both imperial and local history, profoundly reshaping the region's social, economic, and political structures[6].

Roman provinces functioned as administrative units governed by officials appointed by the emperor or Senate, serving as mechanisms for the diffusion of Roman law, infrastructure, and culture, while also facilitating the extraction of local resources[2]. The establishment of the province of Dacia was driven by both strategic and economic considerations. Its abundant natural resources – particularly the gold and silver mines of Transylvania – represented a major financial asset for the Empire[3], while the conquest neutralized a persistent military threat along the Danube frontier[4].

Despite the importance of these events, historiographical sources for the Dacian Wars remain limited. Knowledge of the conflict relies primarily on the account of Cassius Dio[5], fragments attributed to Criton [6], a fragment of Trajan's *commentarii* on the Dacian Wars and monumental visual narratives such as Trajan's Column in Rome[7,8], and the *Tropaeum Traiani* in Moesia Inferior[9].

Trajan's interest in conquering Dacia was multifaceted. Beyond the economic and strategic benefits, the conquest served to enhance Trajan's prestige as a military commander and solidify his authority as emperor. The influx of gold from Dacia's mines helped funding numerous public monuments and infrastructure projects across the Empire[10]. The conquest also provided a strategic advantage, enabling better protection and defense of the Roman Empire's Danube frontier[11,12], while also serving financial aims, including the cessation of subsidy payments established during Domitian's reign[12,13].

The military campaign against Dacia was a formidable undertaking, involving the deployment of nearly half of the Roman legions[14]. Trajan's preparation for the war included the construction of roads and alliances with neighboring tribes to isolate the Dacian king, Decebalus. The Roman forces, totaling more than 150,000 soldiers, faced a Dacian army familiar with the terrain and supported by local alliances[15]. Despite this, the conflict culminated in the siege of Sarmizegetusa Regia[16], the Dacian capital, and the eventual suicide of Decebalus, which marked the end of Dacian resistance[17]. The famous visual representation of the war on Trajan's column corroborates the literary tradition of captivity and enslavement of Dacian prisoners on a large scale[18]. Conscription into Roman auxiliary units also took many more Dacian men of fighting age away from their homeland.

The spoils of war, including the vast Dacian treasury, played a crucial role in enhancing Rome's stock of precious metals[3]. Trajan's conquest of Dacia not only legitimized his authority but also strengthened Rome’s economic and military position. The province of Dacia, strategically organized to exploit local resources, became a vital part of the Roman Empire, with a governor of consular rank overseeing its administration[2].

The Roman rule in Dacia led to significant social and cultural changes, with the local Dacian elite effectively dismantled as a consequence of death and captivity in warfare, and replaced by Roman administrative structures[19]. The province saw extensive colonization, both military and civilian, with settlers from across the Roman world taking advantage of economic opportunities. Although Roman occupation had a profound impact, biological evidence for such transformative changes remains elusive in both bioarcheological and historical contexts. This intense colonization marked the aftermath of Dacia’s conquest, with the most notable example being the founding of the colony of Ulpia Traiana Sarmizegetusa between 108 and 110 CE, located approximately 40 km from the former Dacian capital, Sarmizegetusa Regia[20]. This new Roman city served as the administrative and political center of the province and a prominent expression of Roman culture. The consular governor resided in Ulpia Traiana Sarmizegetusa when exercising his jurisdiction, while also maintaining a presence in Apulum[21], the base of Legio XIII Gemina[22], where he acted as the military commander. This dual role underscores the strategic importance of both locations in the governance and defense of Roman Dacia.

The initial Roman occupation phase involved the establishment of military camps, where civilians lived alongside the Roman army, fostering trade and improving local livelihood[12]. These camps required constant supplies for food, clothing, weapons, tools, and other essentials, creating economic opportunities for local farmers, artisans, and traders, who began to depend economically on the presence of Roman soldiers. During campaigns, civilians followed troops, leading to the creation of settlements called canabae (outside legionary fortresses)[23] and vici (outside auxiliary forts)[24], which facilitated the spread of Roman influence in the new province. Civilian settlements developed just outside Roman military camps, housing traders, craftsmen, soldiers’ families, and local laborers, eventually growing into thriving commercial hubs, although marriage restrictions for soldiers persisted until the reign of Emperor Septimius Severus[25]. Moreover, Roman veterans settled in areas where they served, particularly in Roman Dacia, contributing agricultural expertise and enhancing living standards through economic opportunities[12].

After Trajan’s death in 117 CE, his successor emperor Hadrian sought to consolidate the Empire’s borders[26], recognizing that the newly conquered territories – such as Dacia – were too demanding to control[27]. This decision was influenced by the military challenges in the Danubian region, where the Roman army faced considerable difficulties in dealing with uprisings from barbarian populations, such as the Sarmatians, nomadic Iranian-speaking groups who inhabited a vast region extending from the Pontic-Caspian steppe to the Carpathian Mountains[28], and whose raids even led to the death of the former governor of Dacia, Gaius Julius Quadratus Bassus[29].

As a consequence, Hadrian reorganized the province into two independent entities: Dacia Inferior, which included the territories previously assigned to Lower Moesia, excluding the plains of current

Muntenia and the south of Moldova, which were ceded to the Sarmatian Roxolani, and Dacia Superior, which corresponded to the old Province of Dacia[30]. Shortly afterwards, towards the end of 119 CE, a third province was established in the north, Dacia Porolissensis, with its capital at Porolissum[31], to facilitate defense against the northwest, where Dacian tribes and significant Germanic groups resided[32]. Dacia Superior (Upper Dacia) oversaw the resources of the Empire and controlled the population of the Sarmatian Iazyges, while Dacia Inferior (Lower Dacia) monitored the movements of the nomadic Sarmatian Roxolani and Dacian populations of the Muntenia region. Dacia Superior legion, Legio XIII Gemina, stationed in Apulum and was governed by a praetorian governor, while the other two provinces – Dacia Porolissensis and Dacia Inferior – had equestrian governors and auxiliary troops. These three Dacias were not interconnected, and each had its own governor with full authority within the province, with a very elusive form of coordination[33]. Dacia Porolissensis retained its capital at Porolissum, Dacia Superior was renamed Dacia Apulensis, with its capital at Apulum, and Dacia Inferior became Dacia Malvensis, with its capital at Malva. This organization persisted until the reign of Gallienus – the Roman emperor during the turbulent Crisis of the Third Century[34], from 253 to 268 CE – and the eventual abandonment of the province under Emperor Aurelian around 271 CE. The consequences of the Marcomannic Wars – a series of conflicts fought under Marcus Aurelius between 166 and 180 between the Roman Empire and a coalition of Germanic and Sarmatian tribes – limited the development of Roman Dacia and led to a decrease in economic activities[35].

During the 50 years of military anarchy in the 3rd century CE, Roman Dacia lost its fundamental role as it could no longer prevent the entry of barbarian populations into the Danube area. Consequently, Emperor Aurelian, in 271 CE, decided to withdraw Roman troops from Dacia[4]. Subsequent Emperors, such as Diocletian, paid significant attention to the refortification of the northern bank of the Danube[36], while Constantine the Great's relocation of the capital to Constantinople (modern-day Istanbul) made the Danubian region more strategically important[37]. The construction in 328 CE of a bridge over the Danube linking Sucidava and Colonia Ulpia Oescus, during the reign of Constantine the Great, reflects continued Roman strategic interest in the region. Thus, the political influence of the Roman state remained significant even after the formal abandonment of the province of Roman Dacia.

#### References


1. Breeze, D. J. *Frontiers of the Roman Empire: Hadrian's Wall: Der Hadrianswall / Le Mur d'Hadrien*. (Archaeopress, 2023).

2. Burnett, A. The Roman Provinces, 300 BCE–300 CE: Using Coins as Sources. *Higher Education from Cambridge University Press* https://www.cambridge.org/highereducation/books/the-roman-provinces-300-bce300-ce/4DAACB2DADBDDCB78ED19901723D4B03 (2024) doi:10.1017/9781009420099.

3. Green, G. A. & Smythe, D. Tracing Dacian gold in Roman *aurei*. *J. Archaeol. Sci. Rep.* **39**, 103128 (2021).

4. Ellis, L. 'Terra Deserta': Population, Politics, and the [de]Colonization of Dacia. *World Archaeol.* **30**, 220–237 (1998).

5. Wheeler, E. L. Roman's Dacian Wars: Domitian, Trajan, and Strategy on the Danube, Part I. https://www.academia.edu/4104242/Romans_Dacian_Wars_Domitian_Trajan_and_Strategy_on_the_Danube_Part_I.

6. Savo, M. B. Tito Statilio Critone: medico letterato e storico delle guerre daciche. *Tradiz. E Trasm. Degli Stor. Greci Fram. Ricordo Silvio Accame Atti II Workshop Internazionale Roma 16-18 Febbraio 2006 Tivoli TORED 2009* https://www.academia.edu/15026150/Tito_Statilio_Critone_medico_letterato_e_storico_delle_guerre_daciche.

7. Poulter, A. G. Trajan's Column and the Dacian Wars. *Britannia* **23**, 331–333 (1992).

8. Fox, A. TRAJANIC TREES: THE DACIAN FOREST ON TRAJAN'S COLUMN. *Pap. Br. Sch. Rome* **87**, 47–69 (2019).

9. Black, E. L. Tropaeum Traiani (Adamklissi). in *The Encyclopedia of Ancient History* (John Wiley & Sons, Ltd, 2012). doi:10.1002/9781444338386.wbeah16158.

10. Griffin, M. Nerva to Hadrian. in *The Cambridge Ancient History, XI. The High Empire, A.D. 70-192* 84–131 (Cambridge University Press, Cambridge, 2000).

11. Salmon, E. T. Trajan's Conquest of Dacia. *Trans. Proc. Am. Philol. Assoc.* **67**, 83–105 (1936).

12. Oltean, I. A. *Dacia: Landscape, Colonization and Romanization*. (Routledge, London, 2007). doi:10.4324/9780203945834.

13. Oltean, I. A. & Hanson, W. S. Conquest strategy and political discourse: new evidence for the conquest of Dacia from LiDAR analysis at Sarmizegetusa Regia. *J. Roman Archaeol.* **30**, 429–446 (2017).

14. Pădureanu, E. D. A possible attack direction used by the Roman army during the Dacian Wars. https://doi.org/10.3406/valah.2013.1136 (2013) doi:10.3406/valah.2013.1136.

15. Sidebottom, H. The Dacian Wars, 84–106. in *The Encyclopedia of Ancient Battles* 1–5 (John Wiley & Sons, Ltd, 2017). doi:10.1002/9781119099000.wbabat0640.

16. Comes, R. *et al.* ENHANCING ACCESSIBILITY TO CULTURAL HERITAGE THROUGH DIGITAL CONTENT AND VIRTUAL REALITY: A CASE STUDY OF THE SARMIZEGETUSA REGIA UNESCO SITE. *J. Anc. Hist. Archaeol.* **7**, (2020).

17. Speidel, M. P. The Suicide of Decebalus on the Tropaeum of Adamklissi. *Rev. Arch□ologique* 75–78 (1971).

18. Hölscher, T. Images of War in Greece and Rome: Between Military Practice, Public Memory, and Cultural Symbolism. *J. Roman Stud.* **93**, 1–17 (2003).

19. Varga, R. THE NATURE OF ROMAN DOMINION OVER THE PROVINCE OF DACIA NOTES ON THE ROMANIZATION PHENOMENON AND ITS LIMITS. *J. Anc. Hist. Archaeol.* **8**, (2021).

20. Alexandrescu, C.-G. Ulpia Traiana Sarmizegetusa, Romania. in *The Encyclopedia of Ancient History* 1–1 (John Wiley & Sons, Ltd, 2022). doi:10.1002/9781444338386.wbeah16160.pub2.

21. Diaconescu, A. Dacia. in *A Companion to the Archaeology of the Roman Empire* 273–296 (John Wiley & Sons, Ltd, 2024). doi:10.1002/9781118538265.ch13.

22. Moga, V. PREFECȚI AI CASTRULUI LEGIUNII XIII GEMINA LA APULUM. *Apulum* **13**, 651–657 (1975).

23. Sorin, N. Canabae legionis in Dacia. Military presence and municipal evolution at Apulum and Potaissa. *ALFRED VON DOMASZEWSKI Lat. Epigr. Roman Emp. Acts Colloq. Held Timișoara Dec. 14th–17th 2022* https://www.academia.edu/122574506/Canabae_legionis_in_Dacia_Military_presence_and_municipal_evolution_at_Apulum_and_Potaissa (2024).

24. Hanel, N. Military Camps, Canabae, and Vici. the Archaeological Evidence. in *A Companion to the Roman Army* 395–416 (John Wiley & Sons, Ltd, 2007). doi:10.1002/9780470996577.ch23.

25. Campbell, B. The Marriage of Soldiers under the Empire. *J. Roman Stud.* **68**, 153–166 (1978).

26. Gardner, A. Hadrian's Wall and Border Studies: Problems and Prospects. *Britannia* **53**, 159–171 (2022).

27. Ardevan, R. & Zerbini, L. *La Dacia romana - Rubbettino editore*. (Rubettino, 2007).

28. Fischer, T. Archaeological Evidence of the Marcomannic Wars of Marcus Aurelius (AD 166–80). in *A Companion to Marcus Aurelius* 29–44 (John Wiley & Sons, Ltd, 2012). doi:10.1002/9781118219836.ch2.

29. Ethnicity and Identity in the Roman Empire. in *Rome: An Empire of Many Nations: New Perspectives on Ethnic Diversity and Cultural Identity* (eds Price, J. J., Finkelberg, M. & Shahar, Y.) 15–84 (Cambridge University Press, Cambridge, 2021).

30. Syme, R. Review of Die Reichsbeamten von Dazien. *J. Roman Stud.* **36**, 159–168 (1946).

31. De Sena, E. C. Porolissum. in *The Encyclopedia of Ancient History* (John Wiley & Sons, Ltd, 2012). doi:10.1002/9781444338386.wbeah16110.

32. Wade, D. W. More Ado about Dacia. *Class. World* **64**, 114–116 (1970).

33. Potter, D. Procurators in Asia and Dacia under Marcus Aurelius: A Case Study of Imperial Initiative in Government. *Z. Für Papyrol. Epigr.* **123**, 270–274 (1998).

34. Hartmann, U. The Third-Century "Crisis". in *The Encyclopedia of Ancient Battles* 1–22 (John Wiley & Sons, Ltd, 2017). doi:10.1002/9781119099000.wbabat0720.

35. Ehrhardt, C. What Should One Do about Dacia? *Class. World* **63**, 222–226 (1970).

36. Bird, H. W. Diocletian and the Deaths of Carus, Numerian and Carinus. *Latomus* **35**, 123–132 (1976).

37. Gandila, A. Fighting against nature: Romans and Barbarians on the Icy Danube. https://philpapers.org/rec/GANFAN-2 (2022).
