## Supplementary figures and images for "On the Edge of Empire: Paleogenomic Insights into Roman Dacia"

### Supplemental Figure S1

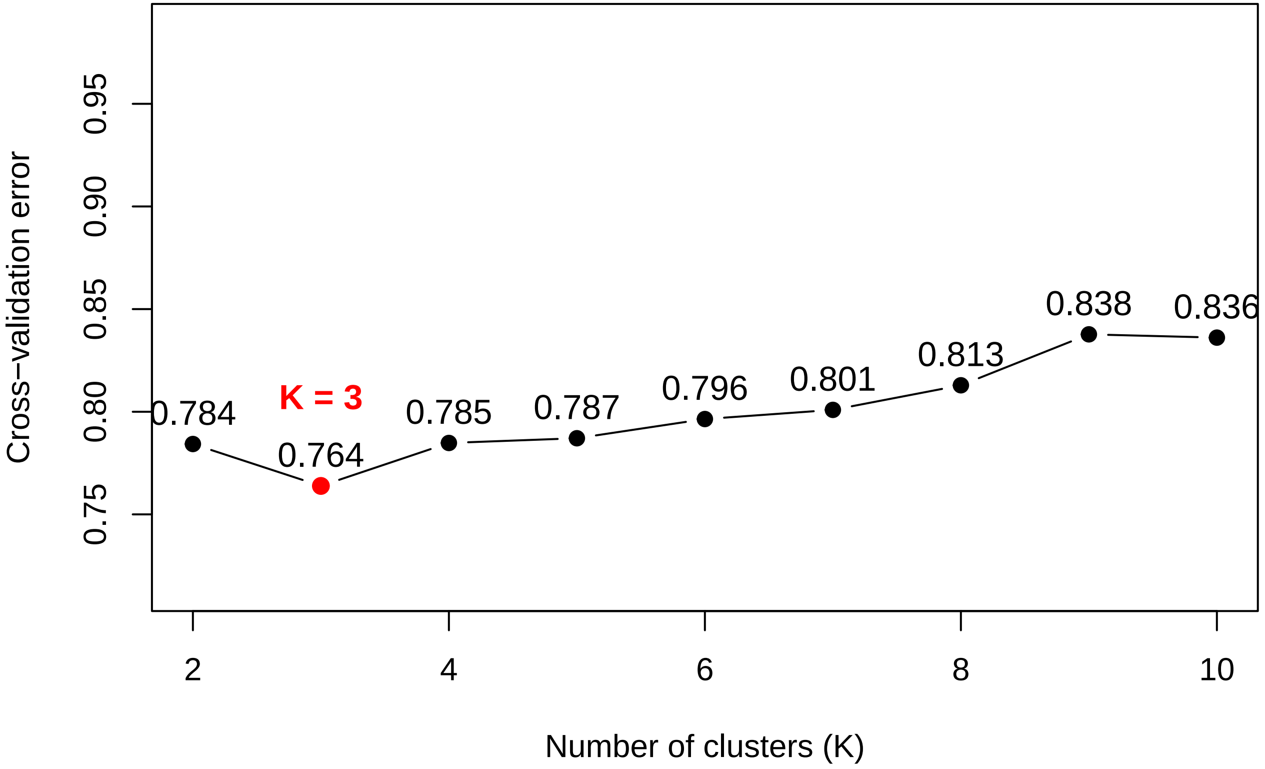
